# Isolation and Characterization of Extracellular Vesicles from the Fungal Phytopathogen *Colletotrichum higginsianum*

**DOI:** 10.1101/2022.01.07.475419

**Authors:** Brian D. Rutter, Thi-Thu-Huyen Chu, Kamil K. Zajt, Jean-Félix Dallery, Richard J. O’Connell, Roger W. Innes

## Abstract

Fungal phytopathogens secrete extracellular vesicles (EVs) associated with enzymes and phytotoxic metabolites. While these vesicles are thought to promote infection, defining the true contents and functions of fungal EVs, as well as suitable protein markers, is an ongoing process. To expand our understanding of fungal EVs and their possible roles during infection, we purified EVs from the hemibiotrophic phytopathogen *Colletotrichum higginsianum*, the causative agent of anthracnose disease in multiple plant species, including *Arabidopsis thaliana*. EVs were purified in large numbers from the supernatant of protoplasts but not the supernatant of intact mycelial cultures. We purified two separate populations of EVs, each associated with over 700 detected proteins, including proteins involved in vesicle transport, cell wall biogenesis and the synthesis of secondary metabolites. We selected two SNARE proteins (Snc1 and Sso2) and one 14-3-3 protein (Bmh1) as potential EV markers and generated transgenic lines expressing fluorescent fusions. Each marker was confirmed to be protected inside EVs. Fluorescence microscopy was used to examine the localization of each marker during infection on *Arabidopsis* leaves. These findings further our understanding of EVs in fungal phytopathogens and will help build an experimental system to study EV inter-kingdom communication between plants and fungi.

## Introduction

Secretory pathways are essential to the life of a fungus. They contribute to the synthesis and remodeling of cell walls, nutrient acquisition and host/symbiont interactions (Krijger et al., 2014; Latge, 2007; McCotter et al., 2016). In particular, plant fungal pathogens rely heavily on secreted proteins to invade and manipulate their hosts. Compared to saprophytes and animal pathogens, phytopathogens have a considerably larger secretome to meet the demands of their vegetarian lifestyles (Krijger et al., 2014). Phytopathogens secrete an arsenal of cell wall-degrading enzymes that allow them to penetrate deep into plant tissues and break down surrounding materials to release nutrients (Kubicek et al., 2014). In addition, a subset of secreted protein effectors help phytopathogens evade detection by the plant immune system by masking chitin molecules that would otherwise be recognized by receptors on the plant cell membrane (de Jonge et al., 2010; van Esse et al., 2007). Other secreted proteins neutralize plant-produced antimicrobial enzymes, such as chitinases, proteases and peroxidases through inhibition or modifications (Hemetsberger et al., 2012; Jashni et al., 2015; Naumann & Price, 2012; Naumann & Wicklow, 2010; Naumann et al., 2009; Naumann et al., 2011; van Esse et al., 2008). Phytopathogens also secrete a wide array of small effector proteins that function in the apoplast or host cytoplasm to suppress plant immune responses (Pradhan et al., 2021). Finally, secondary metabolites released by phytopathogens can damage host tissues or interfere with plant defenses and associated signaling pathways (Collemare et al., 2019; Peng et al., 2010).

Most extracellular proteins are released through the conventional secretory pathway, in which proteins possessing an N-terminal signal peptide are co-translationally imported into the lumen of the endoplasmic reticulum (ER). From the ER, proteins are passed to the Golgi apparatus and in the process become modified through cleavage, additional folding and glycosylation. From the Golgi apparatus, proteins are transported in secretory vesicles to the plasma membrane (PM) for secretion (Peberdy, 1994; Shoji et al., 2014). While the conventional pathway is the best studied route for secreted proteins, there are alternative, unconventional pathways. Proteins can be secreted directly across the PM through pores or via ABC-transporters, and some secretory vesicles released from the ER can bypass the Golgi apparatus completely (Rabouille, 2017). Proteins can also be released through a vesicular mechanism involving the production of extracellular vesicles (EVs) (Rodrigues et al., 2013; Shoji et al., 2014).

EVs are nano-sized, lipid-bilayer compartments that function in the extracellular transport of proteins, lipids, nucleic acids and other macromolecules. EVs are generated through two main processes: 1) they can pinch off directly from the PM or 2) they can form as intraluminal vesicles inside late endosomes (multivesicular bodies) and are then released upon fusion of the late endosome with the PM (van Niel et al., 2018). Although the majority of EV research has been conducted in mammalian systems, the phenomenon of EV secretion has been observed in all domains of life (Woith et al., 2019; Yanez-Mo et al., 2015).

Fungal EVs were first isolated in 2007 from the opportunistic human pathogen *Cryptococcus neoformans* and have since been isolated from multiple species of yeasts and filamentous fungi (Rizzo et al., 2020; Rodrigues et al., 2007). Fungal EVs contain a diverse cargo of lipids, polysaccharides and secondary metabolites. They are enriched for proteins involved in cellular metabolism, translation, transport, signaling and stress responses (Bleackley, Dawson, et al., 2019). Similar to mammalian EVs, fungal EVs also contain a variety of RNA molecules, including small, non-coding and messenger RNAs (Peres da Silva et al., 2015).

Functionally, fungal EVs have been described as “virulence bags” containing multiple factors important for pathology (Albuquerque et al., 2008; Matos Baltazar et al., 2016; Rodrigues et al., 2007; Vallejo et al., 2012; Vargas et al., 2015). Injections of fungal EVs in mice have been shown to enhance the virulence of fungal pathogens and promote the colonization of organs (Huang et al., 2012; Ikeda et al., 2018; Marina et al., 2020). Fungal EVs also contribute to pathology through cell-to-cell communication by promoting biofilm formation and antibiotic resistance, as well as by inducing heightened virulence states in EV-treated fungal cells (Hai et al., 2020; Zarnowski et al., 2018)

Fungal EVs have also been linked to cell wall synthesis. Under conditions that stimulate the regeneration of the cell wall, protoplasts of *Aspergillus fumigatus* secrete high numbers of EVs. These EVs are associated with carbohydrates and proteins essential for the synthesis of major cell wall components (ex: β-1,3- and α-1,3-glucans, chitin, galactomannan and galactosaminogalactan) (Rizzo et al., 2020). EVs of *Saccharomyces cerevisiae* are also associated with glucan and chitin synthases. The presence of these proteins as EV cargo allows the vesicles to rescue yeast cells treated with an inhibitor of β-1,3-glucan synthesis (Zhao et al., 2019). It has been suggested that the cell wall remodeling capabilities of EVs facilitate their passage through the fungal cell wall to reach the extracellular space, but this has not yet been fully verified (Rizzo et al., 2020).

While the majority of fungal EV studies have focused on human pathogens, there is a growing body of research on plant pathogens. In fact, the first example of EVs isolated from a filamentous fungus came from *Alternaria infectoria*, the causative agent of black point and leaf blight diseases in grain crops (Silva et al., 2014). More recent studies have examined EVs from *Zymoseptoria tritici* (septoria leaf blotch in wheat), *Fusarium oxysporum f. sp. vasinfectum* (*Fusarium* wilt disease in cotton), *Ustilago maydis* (maize smut) and *Fusarium graminearum* (*Fusarium* stalk rot) (Bleackley, Samuel, et al., 2019; Garcia-Ceron, Dawson, et al., 2021; Garcia-Ceron, Lowe, et al., 2021; Hill & Solomon, 2020). EVs from these phytopathogens resemble other fungal EVs in their size and contents. They are enriched for proteins involved in primary cellular metabolism and can contain full length mRNAs (Bleackley, Samuel, et al., 2019; Hill & Solomon, 2020; Silva et al., 2014). EVs from *F. oxysporum f. sp. vasinfectum* also co-purify on a density gradient with a phytotoxic, purple-colored metabolite that causes tissue collapse when injected into leaves of cotton plants or *Nicotiana benthamiana* (Bleackley, Samuel, et al., 2019). In all cases, the EVs from fungal phytopathogens are postulated to promote virulence and may use metabolic enzymes to interfere or even reprogram host plant cells (Bleackley, Samuel, et al., 2019; Kwon et al., 2021; Silva et al., 2014).

While research into fungal EVs is growing, questions regarding their biogenesis, ultimate functions and ability to cross the fungal cell wall remain unanswered. Researchers have also struggled to define fungal EV biomarkers. Tetraspanin proteins such as CD63 are widely used as EV markers in mammalian systems (Choi et al., 2015). While lacking true orthologs to human tetraspanins, filamentous fungi do encode proteins having a very similar structure, namely four transmembrane domains, an intracellular loop and two extracellular loops, the larger of which contains the highly conserved CCG cysteine pattern that is a hallmark of mammalian tetraspanins (Lambou et al., 2008). However, these fungal tetraspanins have so far not been detected in any fungal EV proteomes. Similarly, although fungi possess orthologs to components of the ESCRT complex, another common mammalian EV marker, they are often present in low abundance or not detectable in fungal EVs (Bleackley, Dawson, et al., 2019). A further problem is that fungal EVs are rarely density-gradient purified. This means that the majority of published fungal EV proteomes, sources for selecting biomarkers, likely contain co-pelleting, extra-vesicular molecules.

In this study, we expand the current knowledge of fungal EVs by isolating EVs from the phytopathogen *Colletotrichum higginsianum*. Members of the genus *Colletotrichum* are mostly hemibiotrophs that cause anthracnose leaf spot diseases on thousands of plant species and are mostly hemibiotrophic pathogens, meaning that they initially invade and derive nutrients from living plant cells (biotrophy) before later switching to a destructive necrotrophic phase, when they feed on dead host cells (O’Connell et al., 2012). *C. higginsianum* is adapted to infect cruciferous plants, including not only crop species but also the model plant *Arabidopsis thaliana.* The pathosystem formed between these two organisms is regarded as a model for hemibiotrophic plant-fungus interactions (O’Connell et al., 2004). Here, we report that *C. higginsianum* is also capable of secreting EVs. These EVs are not readily released from vegetative mycelia grown in liquid media, but they can be isolated in large numbers after removal of the cell wall. Two separate populations of EVs were purified from *C. higginsianum* protoplasts based on their buoyant density. Vesicles were treated with protease to remove extra-vesicular contaminants and then examined for their protein contents. We report several hundred proteins, including those involved in cell wall synthesis/remodeling and the production of secondary metabolites. We further developed transgenic, fluorescent EV biomarker lines that could be used in the future to track EV secretion during infection. This research will strengthen fungal EV research and lay the groundwork for modeling inter-kingdom EV communication in plant-fungal interactions.

## Materials & Methods

### Fungal material

*Colletotrichum higginsianum* isolate IMI349063A was used for all experiments and as a background strain for the generation of transgenic lines. Stocks of fungal spores were stored in 15% glycerol and 1X potato dextrose broth at −80°C. Cultures were prepared fresh from stock and grown on 100 ml of Mathur’s agar (2.8 g/L glucose, 2.2g Mycological peptone, 1.2 g/L MgSO_4_ × 7H_2_O, 2.7g/L KH_2_PO_4_, 30 g/L agar, pH 5.5) in 250 ml flasks. To create liquid cultures, spores from 7-14-day old cultures were collected from the surface of solid media into sterile, deionized water. Spores were inoculated at a final concentration of 5 × 10^5^ spores/ml in liquid Mathur’s medium. 200-500 ml liquid cultures were grown in the dark at 25°C and 100 rpm for 3 days.

### Transmission electron microscopy (TEM)

Hypocotyl segments (7 cm) were excised from 6-day-old seedlings of *Phaseolus vulgaris* cv. Kievitsboon KoeKoek and the cut ends were sealed with paraffin wax. The segments were placed horizontally on glass supports over wet paper inside plastic boxes, then inoculated with droplets (7 μL) of *C. lindemuthianum* spore suspension (5 × 10^5^ spores/ml) and incubated at 17 °C. At 4 days after inoculation, strips of hypocotyl tissue (c. 0.5 mm thick and 5 mm long) were cut parallel to the hypocotyl surface with a razor blade. Infected areas were excised using a 2 mm diameter biopsy punch and vacuum-infiltrated with 1-hexadecene, before mounting in 1-hexadecene between two aluminium sample holders (0.3 mm deep) and cryo-fixing in a Balzers HPM 010 high-pressure freezer. Tissues were freeze-substituted with acetone containing 2% (w/v) osmium tetroxide at −90 °C for 48 h, −60 °C for 12 h, and −30 °C for 8 h using a Reichert CS freeze-substitution device. Samples were then infiltrated with Epon-Araldite resin and polymerized at 60 °C. Ultrathin sections were mounted on Formvar-coated slot grids, stained with uranyl acetate and lead citrate, and viewed with a Hitachi H7000 TEM. Biotrophic hyphae of *C. higginsianum* were isolated from infected *A. thaliana* leaves by fluorescence-activated cell sorting, cryo-fixed and prepared for TEM as described previously (Takahara et al., 2009).

For negative staining, vesicles resuspended in 20 mM Tris-HCl pH 7.5 were applied to glow-discharged 3.05-mm copper Formvar-carbon-coated electron microscopy grids (Electron Microscopy Sciences). After 1 minute, samples were wicked off using filter paper, and the grids were washed with 2% uranyl acetate. The grids were allowed to air dry and imaged at 80 kV using a JEM-1010 transmission electron microscope (JEOL USA).

### Protoplast generation

For protoplast generation, mycelia from 3-day old liquid cultures were harvested by straining the culture through Miracloth (Millipore Sigma catalog number 475855). The mycelia were transferred to a 50 ml Falcon tube and washed twice in 0.7 M NaCl by pelleting mycelia at 8,000 rpm and 25°C for 5 minutes using a JA25.50 rotor and resuspending in sterile saline solution. After the final wash, mycelia were added to a cell wall digestion solution (0.7 M NaCl, 10 mM phosphate buffer, 100 mg/ml VinoTaste® Pro (Novozymes), pH 5.5) and allowed to acclimate for 10 minutes with gentle rocking. The fungi and cell wall digestion solution were then added to a 100 ml flask and incubated for 3 hours at 25°C and 80 rpm. After digestion, protoplasts were pelleted at 2,000 rpm and 4°C for 8 minutes using a JA25.50 rotor. The supernatant was transferred to a fresh 50 ml Falcon tube and the pelleting step was repeated to remove any residual protoplasts. The supernatant was then filtered through a 0.22 µm filter and kept on ice. Pelleted protoplasts were resuspended in cold, sterile 0.7 M NaCl. Their viability was assessed by mixing with Evans Blue to a final concentration of 0.01% (w/v) and examining protoplasts for dye uptake using a light microscope.

### Crude EV isolation

EVs were isolated from the supernatant of cell wall digestion reactions using differential ultracentrifugation. After filtration, 12 ml samples were added to 14 × 89 mm ultra-clear centrifuge tubes (Beckman Coulter, Inc., Brea, CA) and centrifuged at 10,000g and 4°C for 30 minutes (SW41 rotor, Optima XPN-100 ultracentrifuge; Beckman Coulter, Inc.) to remove any potential large debris. The majority of the supernatant (11.5 ml) was transferred to a fresh tube using a pipette, the volume was brought up to 12 ml using fresh 0.7 M NaCl and the EVs were pelleted at 60,000g and 4°C for 90 minutes (SW41, Optima XPN-100 ultracentrifuge). After this step, 11.5 ml of supernatant were removed using a pipette, and the pellet was resuspended in the remaining 0.5 ml. The EV sample was then transferred to a smaller 13 × 51 mm polycarbonate ultracentrifugation tube (Beckman Coulter, Inc., Brea, CA), brought up to 3 ml using sterile 20 mM Tris-HCl pH 7.5 and pelleted at 40,000g and 4°C for 60 minutes (TLA100.3 rotor, Optima Max-XP ultracentrifuge, Beckman Coulter, Inc.). Finally, the supernatant was completely decanted, the upper 3/4 of the tubes were dried with a Kim Wipe to remove residual liquid.

### Density Purification

EVs were further purified on a discontinuous iodixanol gradient (Optiprep™ Sigma-Aldrich) consisting of 40% (v/v), 20% (v/v), 10% (v/v), and 5% (v/v) iodixanol layers. The EV pellets were resuspended in 1 ml of 20 mM Tris-HCl pH 7.5 and mixed with 2 ml of 60% Optiprep™ (Sigma-Aldrich, St. Louis, MO) to create a 40% layer. The remaining layers were created by diluting a 60% OptiPrep stock solution in 20 mM Tris-HCl pH 7.5. The gradient was formed by sequentially adding each 3 ml layer to a 14 × 89 mm ultra-clear centrifuge tube (Beckman Coulter, Inc.), starting with the 40% solution and ending with the 5% solution. The gradient was centrifuged for 17 hours at 100,000g and 4°C (SW41, Optima XPN-100 ultracentrifuge). After centrifugation, the top 3 ml were discarded. The next six 1 ml fractions were collected for further analysis. Samples were added to 13 × 51 mm polycarbonate ultracentrifugation tubes (Beckman Coulter, Inc., Brea, CA) and brought up to a volume of 3.5 ml using cold, sterile 20 mM Tris-HCl pH 7.5. Samples were centrifuged for 1 hour at 100,000g and 4°C (TLA100.3, Optima Max-XP ultracentrifuge). The supernatant was completely decanted, the upper 3/4 of the tubes were dried with a Kim wipe to remove residual liquid and the pellet was resuspended in 20 mM Tris-HCl pH 7.5.

### NanoParticle Tracking Analysis

EV sample particle concentrations were determined using a ZetaView® Particle Tracking Analyzer (Particle Metrix, Diessen) and associated software. The machine was calibrated using 100 nm polystyrene beads, and samples were diluted in 1 ml of 20 mM Tris-HCl, pH 7.5. The machine operated under the following settings: max. diameter: 500 nm, min. diameter: 5 nm, min. brightness: 20, shutter speed: 100.

### Mass spectrometry (MS) Analysis

Vesicle pellets were resuspended in a buffer containing 8M urea and 100 mM ammonium bicarbonate. Disulfide bonds were reduced by incubation for 45 min at 57 °C with a final concentration of 10 mM Tris (2-carboxyethyl) phosphine hydrochloride (Catalog no C4706, Sigma Aldrich). A final concentration of 20 mM iodoacetamide (Catalog no I6125, Sigma Aldrich) was then added to alkylate these side chains and the reaction was allowed to proceed for one hour in the dark at 21 °C. A total of 500 ng trypsin was added (V5113, Promega) and the samples were digested for 14 hours at 37 °C. The following day, the sample was dried down and resuspended in 30 μL 0.1% formic acid, then desalted using a zip tip (EMD Millipore).

Samples were analyzed by LC-MS on an Orbitrap Fusion Lumos (ThermoFisher) equipped with an Easy NanoLC1200 HPLC (ThermoFisher). Peptides were separated on a 75 μm × 15 cm Acclaim PepMap100 separating column (Thermo Scientific) downstream of a 2 cm guard column (Thermo Scientific). Buffer A was 0.1% formic acid in water. Buffer B was 0.1% formic acid in 80% acetonitrile. Peptides were separated on a 120 minute gradient from 4% B to 33% B. Peptides were collisionally fragmented using HCD mode. Precursor ions were measured in the Orbitrap with a resolution of 120,000. Fragment ions were measured in the ion trap. The spray voltage was set at 1.8 kV. Orbitrap MS1 spectra (AGC 4×10^5^) were acquired from 400-2000 m/z followed by data-dependent HCD MS/MS (collision energy 30%, isolation window of 2 Da) for a three second cycle time. Charge state screening was enabled to reject unassigned and singly charged ions. A dynamic exclusion time of 60 seconds was used to discriminate against previously selected ions.

### Database search

The LC-MS/MS data was searched using Proteome Discoverer 2.5 (ThermoFisher Scientific). MS spectra were searched against a *Colletotrichum higginsianum* database downloaded from Uniprot on 06/2018. The database search parameters were set as follows: two missed tryptic cleavage sites were allowed for trypsin digested with 10 ppm precursor mass tolerance and 0.05 Da for fragment ion quantification tolerance. Oxidation of methionine was set as a variable modification. Carbamidomethylation (C; +57Da) was set as a static modification. Results were filtered using the Percolator node with a FDR of 0.01.

### Analysis of protein sequences

Protein sequences were analyzed for predicted signal peptides using SignalP - 5.0 (https://services.healthtech.dtu.dk/service.php?SignalP-5.0, Almagro Armenteros et al., 2019). Transmembrane regions were predicted using TMHMM - 2.0 (https://services.healthtech.dtu.dk/service.php?TMHMM-2.0, Sonnhammer et al., 1998, Krogh et al., 2001). GO term enrichment was determined using FungiFun2 (Priebe et al., 2015).

### Immunoblots

For immunoblots, 40 µl of resuspended vesicles in 20 mM Tris-HCl, pH 7.5 were combined with 10 µl of 5X SDS loading buffer (250 mM Tris-HCl, pH 6.8, 8% SDS, 0.1% Bro-mophenol Blue, 40% glycerol, and 400 mM dithiothreitol) and boiled at 95°C for 5 minutes. Protoplast lysate samples were used as positive controls. Pelleted protoplasts were frozen in liquid nitrogen and ground into a powder using a mortar and pestle. Proteins from the ground protoplasts were extracted in 500 µl of protein extraction buffer (150 mM NaCl, 50 mM Tris HCl, pH 7.5, 0.1% Nonidet P-40, and 1% protease inhibitor cocktail [Sigma-Aldrich, Catalog No.: P2714]) and centrifuged for 5 min at 10,000 rpm and 4°C. Forty microliters of supernatant were combined with 10 µl of 5X SDS loading buffer, and the mixture was boiled at 95°C for 5 minutes. Protein concentrations were determined using Pierce 660nm Protein Assay Reagent (ThermoFisher Scientific). Samples were loaded on 4% to 20% Precise Protein Gels (ThermoFisher Scientific) and separated at 120 V for ~1 h in BupH Tris-HEPES-SDS running buffer (ThermoFisher Scientific). The proteins were transferred to a nitrocellulose membrane (GE Water & Process Technologies). Total protein was visualized with a quick Ponceau S stain. Membranes were washed with 1X Tris-buffered saline containing Tween 20 (TBST; 50 mM Tris-HCl and 150 mM NaCl, pH 7.5, 0.1% Tween 20) and blocked with 5% Difco Skim Milk (BD) overnight at 4°C with gentle rocking. Membranes were incubated with rabbit anti-RFP (Abcam, Catalog No.: ab62341) or anti-mNeonGreen antibody (Chromotek, Catalog No.: 32F6) each at a 1:1,000 dilution or anti-SOD2 (ABclonal, Catalog No.: A1340) at 1:500 for 1 h, washed with TBST, and incubated with horseradish peroxidase-labeled goat anti-rabbit antibody (Abcam, Catalog No.: ab6721) at a 1:5,000 dilution for 1 h. After a final wash in TBST, protein bands were imaged using Immuno-Star Reagents (Bio-Rad) and x-ray film. Band intensities were quantified using ImageJ V2.0.0-rc-43/1.52n.

### Protease Protection Assay

For protease protection assays, crude EV pellets were resuspended in 150 mM Tris-HCl, pH 7.8 and divided into three equal samples. Sample A was treated with buffer only. Sample B was treated with 1 µg/ml trypsin (Promega). Sample C was pre-treated with 1% Triton X-100 (EMD-Millipore) followed by treatment with 1 µg/ml trypsin (Promega). Triton X-100 treatment was carried out on ice for 30 minutes. Trypsin treatment was carried out at 25°C for 1 hour. All samples were subjected to the same incubation temperatures and times.

### Molecular cloning and fungal transformation

Predicted cDNA sequences of putative EV marker genes were amplified from the genomic DNA of *Colletotrichum higginsianum* isolate IMI 349063A including *Ch*Snc1 (protein ID: OBR08408.1, gene ID: CH63R_07173), *Ch*Sso2 (protein ID: OBR07592.1, gene ID: CH63R_09113) and *Ch*Bmh1 (protein ID: OBR15797.1, gene ID: CH63R_00977). All cloning fragments were amplified by PCR using Phusion High-Fidelity DNA polymerase (Thermo Scientific). Plasmids with *Ch*Snc1 and *Ch*Sso2 insertions were made using *in vivo* assembly by homologous recombination in *E. coli*. *ChSnc1* was tagged with the *mScarlet-I* sequence at its 5’ terminus under the control of its native promoter and cloned into the *Sma*I-digested pBIG4MRH binary vector (Tanaka et al., 2007). *Ch*Sso2 was tagged with the *mNeonGreen* sequence at its 5’ terminus under the *ChActin* (Gene ID: CH63R_04240) promoter consisting of 1.7 kb upstream of the start codon; fragments were cloned into *Xmn*I-digested pCGEN plasmid (Motteram et al., 2011). *Ch*Bmh1 was fused to *mCherry* as a C-terminal tag with an intervening linker sequence *ggtcccgggggatcc* and placed under the *ChActin* promoter. The fragments were assembled together with the *Sma*I-digested pBIG4MRH vector using the NEBuilder HiFi DNA Cloning kit according to the manufacturer’s instructions (New England Biolabs). The primers used are summarized in Table S7. Recombinant DNA plasmids were inserted in electrocompetent cells of *Agrobacterium tumefaciens* strain C58C1 then transformed into *C. higginsianum* isolate IMI349063A using *Agrobacterium tumefaciens*-Mediated Transformation (ATMT) (Huser et al., 2009). Transformants of *Ch*Snc1 and *Ch*Bmh1 were selected on potato dextrose agar (PDA) supplemented with hygromycin (100 µg/ml), while *Ch*Sso2 selection was performed on PDA supplemented with geneticin (300 µg/ml).

### Fungal inoculation and confocal microscopy

To determine the subcellular localization of fluorescent fusion proteins in hyphae growing *in vitro*, fungal transgenic lines were grown on Mathur’s agar plates for 3 days at 25°C. Samples were prepared using the ‘inverted agar block method’ (Hickey et al., 2004). A block of agar was cut from the edge of a colony and mounted upside down in a drop of Mathur’s liquid medium on a glass-bottomed Petri dish. For protein localization *in planta*, droplets of fungal spore suspension (5 × 10^5^ spores/ml) were inoculated on cotyledons of 11- to 12-day-old seedlings of *Arabidopsis thaliana* accession Col-0. Seedlings were then placed in a humid box, and incubated at 25°C for the first 24 h in the dark, then with a cycle of 12 h day and 12 h night. Cotyledons were collected at different time points and mounted in perfluorodecalin (Sigma Aldrich) for confocal microscopy (Littlejohn et al., 2014). Observations were made using an inverted confocal microscope (Leica SP5) equipped with a 63x water immersion objective (NA 1.2). For mCherry and mScarlet, both were excited at 561 nm and emission then detected from 600 nm to 630 nm and from 580 to 630 nm, respectively. For mNeonGreen, excitation was at 488 nm and emission was detected from 510 to 540 nm. Fluorescence images were captured in parallel with Differential Interference Contrast (DIC) transmitted light images.

## Results

### EV-like structures are present in the space between the cell wall and plasma membrane in *Colletotrichum* species

Prior to the first isolation of fungal EVs, multiple studies documented the presence of fungal secreted vesicles using transmission electron microscopy. Vesicles have been observed on the cell surface of yeasts and filamentous fungi or sandwiched between the cell wall and plasma membrane, often in a state of budding from the plasma membrane or release from the cell (Anderson et al., 1990; de Paula et al., 2019; Gibson & Peberdy, 1972; Ikeda et al., 2018; Osumi, 1998; Rizzo et al., 2020; Rodrigues et al., 2007; Takeo et al., 1973; Vargas et al., 2015; Wolf et al., 2014). In some yeast species, fungal EVs were seemingly observed in the process of traversing the cell wall (Albuquerque et al., 2008; Anderson et al., 1990; Rodrigues et al., 2007).

To determine whether similar structures could be observed in species of *Colletotrichum*, we used transmission electron microscopy (TEM) to examine the biotrophic hyphae from *C. higginsianum* and *C. lindemuthianum* formed during growth inside living cells of their respective plant hosts (Figure 1). Membrane-bound vesicle-like structures were observed in the paramural space of biotrophic hyphae, between the fungal wall and plasma membrane (Figure 1). These structures varied in size and shape, some appearing spherical, while others were more allantoid-shaped (Figures 1b and 1c). However, from single ultrathin sections it is difficult to tell whether these structures represent discrete vesicles or networks of membranous tubes, similar to those produced by arbuscular mycorrhizal fungi during symbiosis (Ivanov et al., 2019; Roth et al., 2019). The irregular shapes of these structures stand in sharp contrast to the smaller, uniformly spherical intraluminal vesicles observed within nearby fungal multivesicular bodies (MVBs; Figure 1 d). This difference in size and regularity may suggest that the extracellular structures are derived directly from the plasma membrane rather than by the fusion of MVBs with the plasma membrane. Nevertheless, these observations support the presence of EVs in *Colletotrichum* species and, if they are secreted beyond the cell wall as in other fungi, it may be possible to isolate them. We also detected vesicle-like structures in the interfacial matrix between the fungal wall and plant plasma membrane (Figure 1 d), but their fungal or plant origin could not be determined.

**FIGURE 1:**
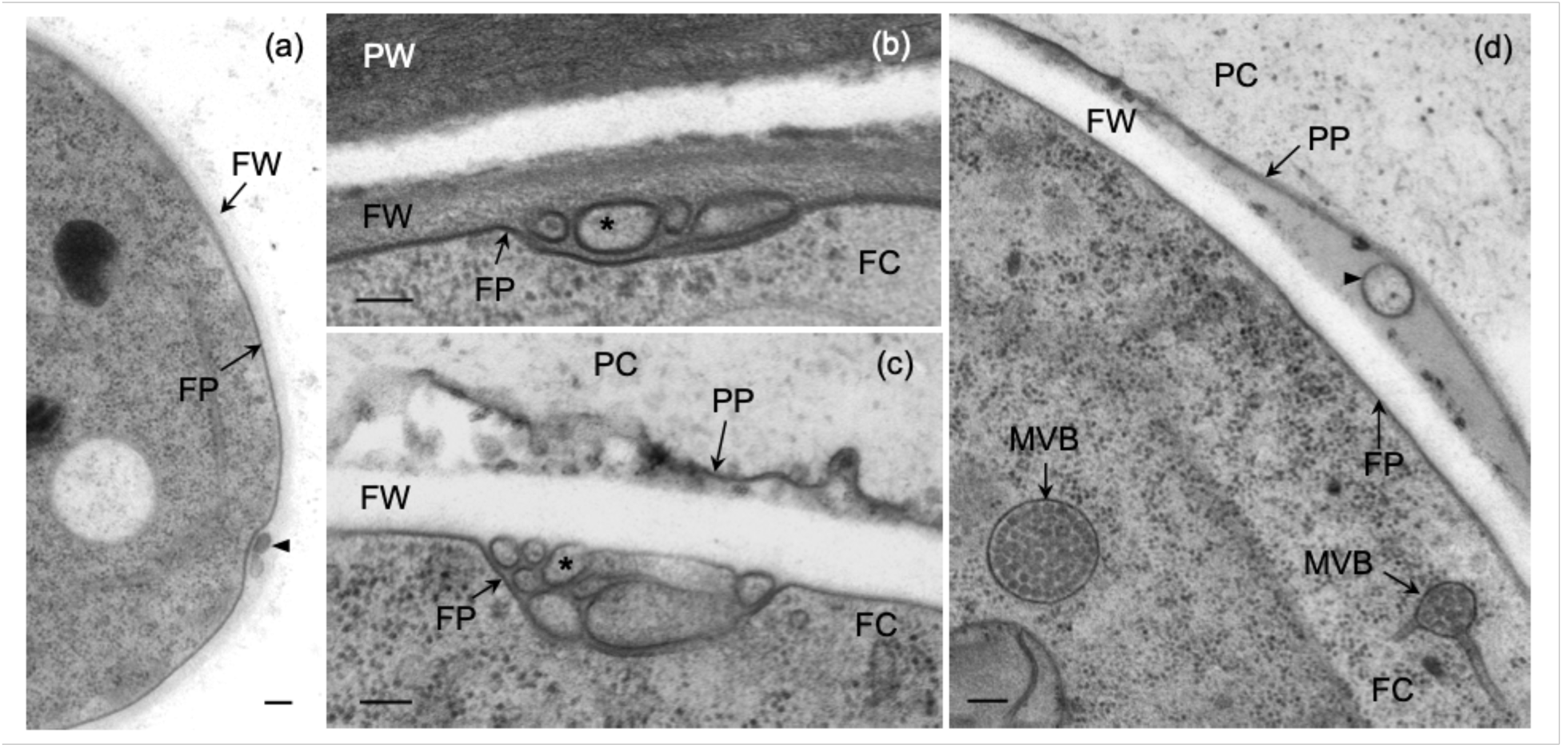
Biotrophic hyphae of *Colletotrichum higginsianum* and *C. lindemuthianum* contain extracellular vesicle-like structures in the paramural space. (a) Biotrophic hypha of *C. higginsianum* isolated from Arabidopsis leaves by fluorescence-activated cell sorting showing paramural vesicles (arrowhead). (b) An extracellular vesicle (arrowhead) inside the interfacial matrix layer between the fungal cell wall (FW) and the plant plasma membrane (PP). (c-d) Biotrophic hyphae of *C. lindemuthianum* infecting *Phaseolus vulgaris* epidermal cells. (c, d) Paramural vesicles (asterisks) between the fungal cell wall (FW) and fungal plasma membrane (FP). All cells were prepared for TEM by high-pressure freezing and freeze-substitution. FC, fungal cytoplasm; MVB, multivesicular body; PC, plant cytoplasm; PW, plant cell wall. Scale bars = 100 nm.

### EV-like particles can be isolated from the supernatant of *C. higginsianum* protoplasts

Fungal EVs were first isolated from the supernatants of *Cryptococcus neoformans* liquid cultures (Rodrigues et al., 2007). This approach has since been used to study EVs in multiple species of yeasts and filamentous fungi, including phytopathogens (Kwon et al., 2021; Rizzo et al., 2020). As a first attempt to isolate EVs from *C. higginsianum*, we processed the supernatant of three-day-old liquid cultures using differential ultracentrifugation. To our surprise, despite processing relatively large volumes of supernatant and using high speeds, we were unable to detect meaningful numbers of EV-like particles. Nano-particle tracking analysis (NTA) failed to identify high numbers of particles in the final pellet (Figure S1 and Figure 3) and while TEM negative staining did reveal some cup-shaped objects reminiscent of vesicles, such structures were rare (Figure S1).

One explanation for the scarcity of EVs in our culture supernatants could be that *C. higginsianum* only secretes EVs under specific circumstances, which we were unable to recreate in the media tested. Another possibility is that *C. higginsianum* EVs were secreted, but the majority remained trapped between the fungal cell wall and plasma membrane. To determine if the cell wall blocked the release of EVs into the liquid culture, we generated *C. higginsianum* protoplasts using a mixture of fungal cell wall-degrading enzymes. Supernatants obtained from protoplast digestion reactions were filtered and processed using differential ultracentrifugation. The resulting crude pellet was then purified on a discontinuous Optiprep gradient.

We selected six fractions from the middle of the density gradient ranging from 10-40% Optiprep (Figure 2a). From our experience with plant EVs, we expect vesicles to float somewhere in this region (Rutter & Innes, 2017). Using NTA, we observed the highest concentrations of particles in fractions three, five and six with corresponding densities of ~1.078, ~1.115 and ~1.173 g/ml. (Figure 2b). Particles in these fractions had an average diameter of ~106, ~102 and ~100 nm, respectively (Figure 2c). Negative staining and TEM analysis revealed abundant, cup-shaped structures in all three fractions (Figure 2d). The size and morphology of the particles in these fractions, as well as their ability to float, suggest they are lipid vesicles. Furthermore, it appears that two separate populations of vesicles were released into the supernatant, a low density (LD) population in fraction three and a high density (HD) population in fractions five and six.

**FIGURE 2:**
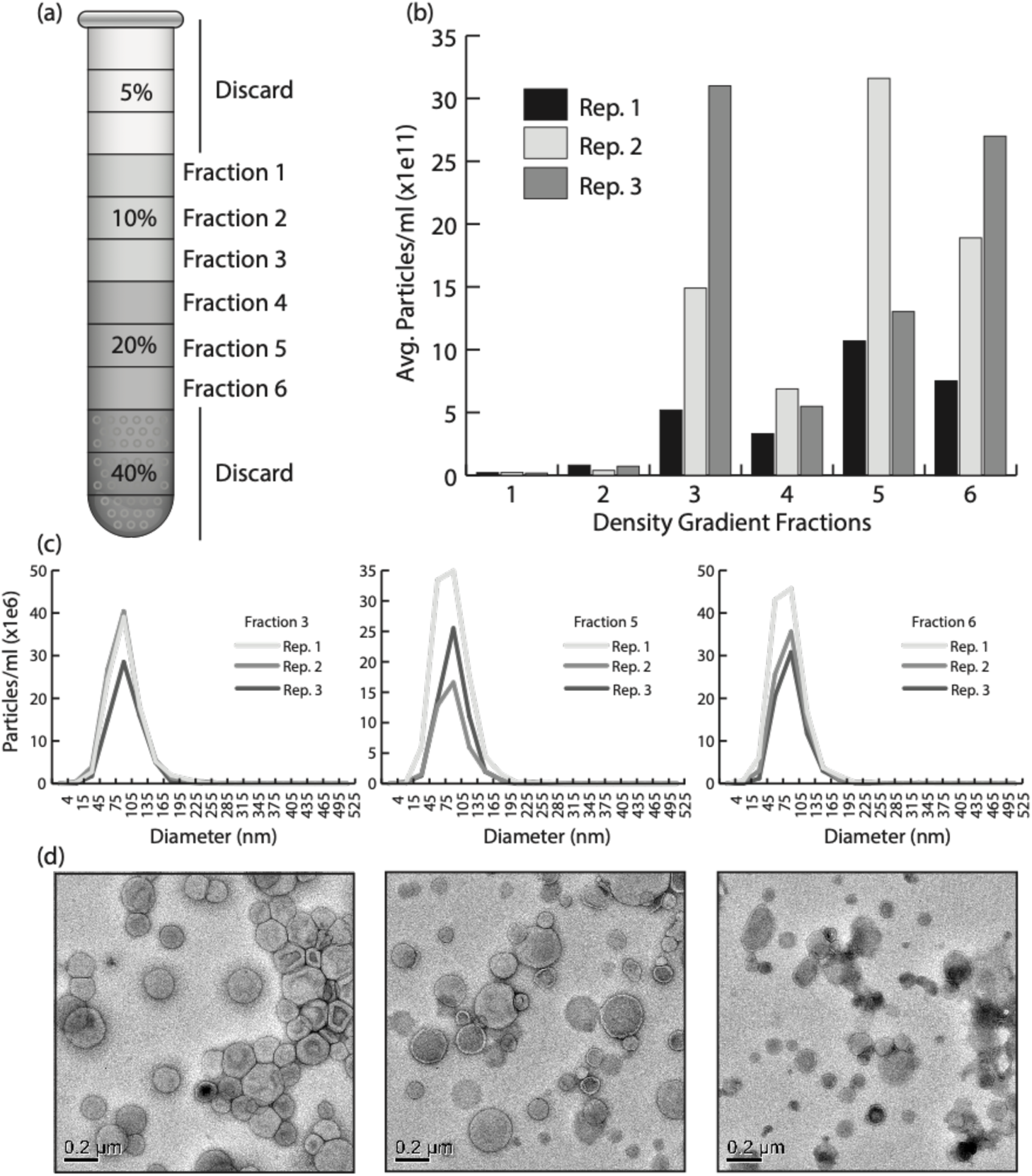
Vesicle-like structures can be purified from *C. higginsianum* protoplasts. (a) Schematic of the Optiprep gradient used to purify vesicles. Crude vesicle pellets were bottom-loaded into a discontinuous Optiprep gradient consisting of 5, 10, 20 and 40% layers. After centrifugation at 100K × g for 17 hrs, the 5% layer was discarded and the next 6 fractions of 1 ml each were collected and processed with further ultracentrifugation to obtain a pure vesicle pellet. (b) Nanoparticle tracking (NTA) data showing the average concentration of particles in each of the collected Optiprep fractions. Three independent experiments are shown in one graph. (c) NTA data showing the size distribution of particles in fractions 3, 5 and 6. Three replicates are shown in each line graph. (d) Transmission electron microscopy negative stain images of fractions 3, 5 and 6.

To demonstrate further that generating protoplasts is key to the isolation of EV-like particles from *C. higginsianum*, we processed the supernatants of undigested mycelia and protoplasts in parallel. Both fungal samples were derived from an equal mass of mycelia and were treated under the same conditions, except that one was incubated in a buffer containing hydrolyzing enzymes and the other was incubated in a buffer lacking enzymes. After differential ultracentrifugation and purification on a density gradient, we detected significantly higher concentrations of particles in fractions isolated from the supernatant of protoplasts compared to those isolated from the supernatant of undigested mycelia (Figure 3). The results show that significant numbers of EV-like particles are only released into the supernatant after the removal of the fungal cell wall. These results also help allay concerns about hyphal tip bursting. Mechanical agitation, as well as changes in osmotic and temperature conditions during the formation of protoplasts can cause the spontaneous bursting of hyphal tips (Bartnicki-Garcia & Lippman, 1972). Such damage would lead to a discharge of intracellular components into the supernatant, which could become confounding artifacts. Our results, however, suggest that if any hyphal tip rupture does occur, it does not generate a significant number of particles. We also assessed the viability of the protoplasts to determine if broken protoplasts might contribute to particle release. Using Evans blue, an exclusion dye that labels cells with damaged plasma membranes (Gaff & Okong’O-Ogola, 1971), we found that the percentage of non-viable protoplasts produced after cell wall digestion was ~1.6% (Figures 4a and 4b). The overwhelming majority of protoplasts (98.4%) remained viable after cell wall digestion, suggesting that the EV-like particles are not derived from damaged protoplasts (Figure 4b).

**FIGURE 3:**
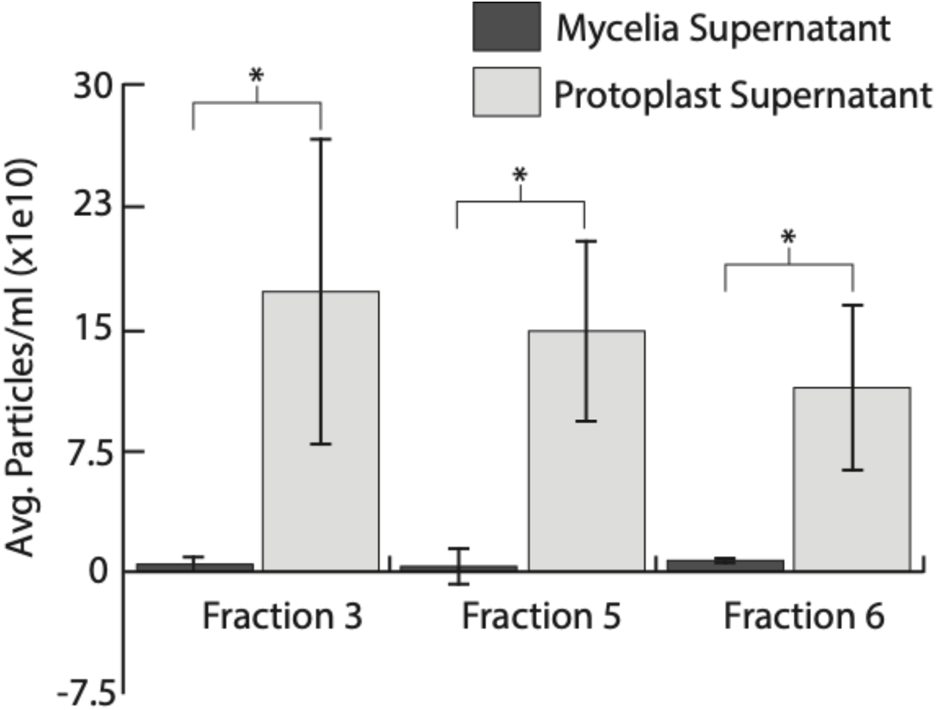
Isolating particles from *C. higginsianum* protoplasts requires removal of the cell wall. Supernatant from mycelia in a mock digest solution or protoplasts were processed in parallel to purify vesicles. Significant numbers of particles were only detected in samples from protoplasts. Bars represent an average of three independent experiments. Error bars represent SD. Asterisks signify a significant difference based on a two-tailed unpaired Student’s *t* test (*P* < 0.05).

**FIGURE 4:**
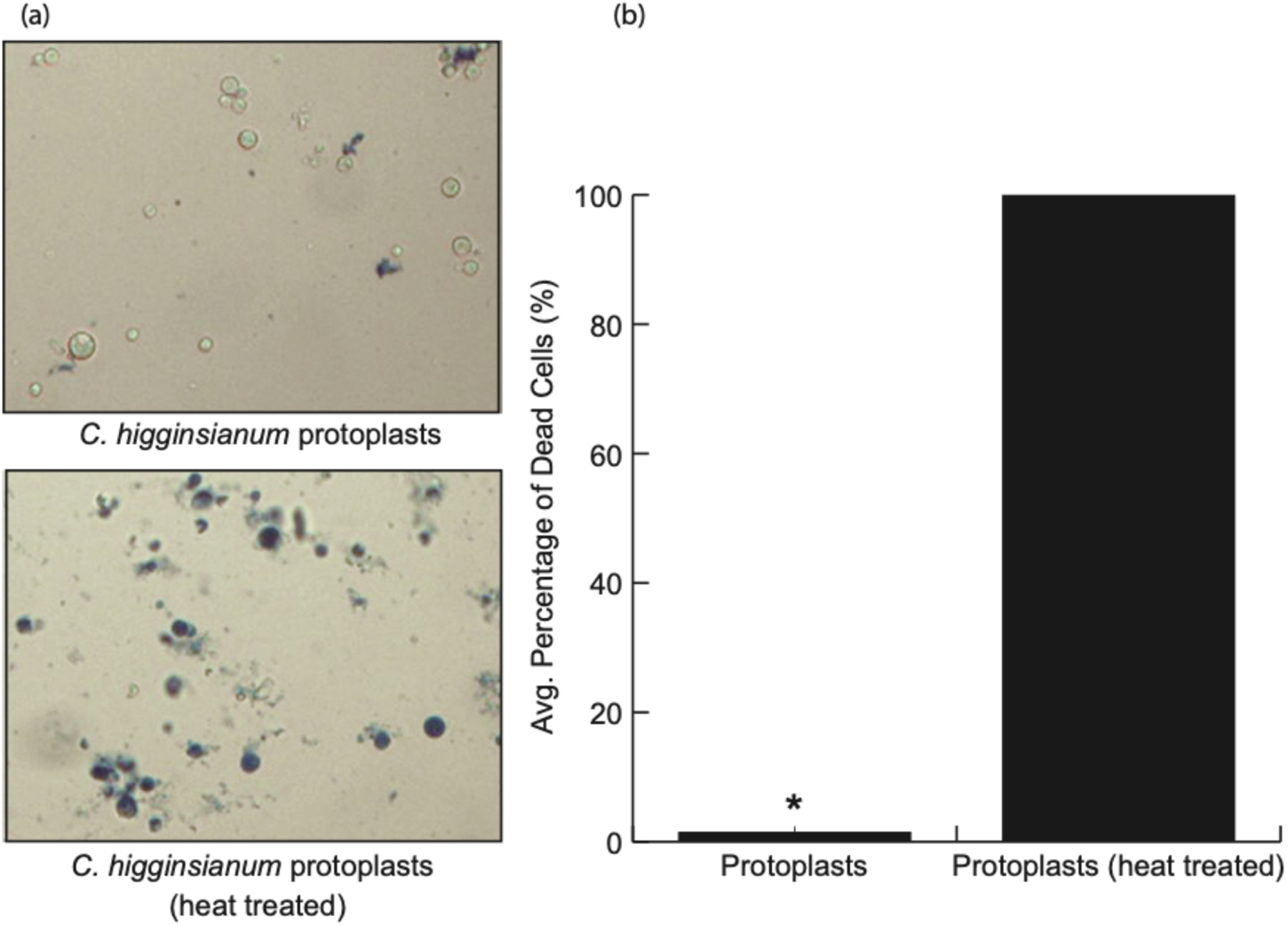
Viability stain of *C. higginsianum* protoplasts. (a) Protoplasts generated after cell wall digestion were mixed with Evans blue dye to a final concentration of 0.04% and observed for staining using a light microscope. A sample of protoplasts was boiled for 5 minutes at 95°C and mixed with dye to provide a positive control for staining. (b) 300 protoplasts were tallied for untreated and heat-treated. The graph represents the mean number of dead protoplasts from three independent experiments. Error bars represent SD. The asterisk signifies the value is significantly different from the other based on a two-tailed unpaired Student’s *t* test (*P* < 1e-8).

### The protein content of *C. higginsianum* EV-like particles

Having purified two populations of *C. higginsianum* vesicles, we next sought to analyze the proteins associated with each population. Crude vesicle samples were treated with trypsin to remove extra-vesicular proteins. The samples were then purified on a density gradient to separate low- and high-density populations and analyzed using LC-MS/MS (Table S1). We selected proteins detected in two out of four replicates with a q-value ≤ 0.01. After applying these criteria, we generated a list of 711 proteins associated with the low-density (LD) vesicles and 707 proteins associated with the high-density (HD) vesicles (Table S2). Approximately 59% of proteins were common to at least two LD replicates and ~64% were common to at least two HD replicates (Figure 5b). The LD and HD populations shared 384 proteins, while 326 were unique to the LD population and 322 were unique to the HD population (Figure 5c).

**FIGURE 5:**
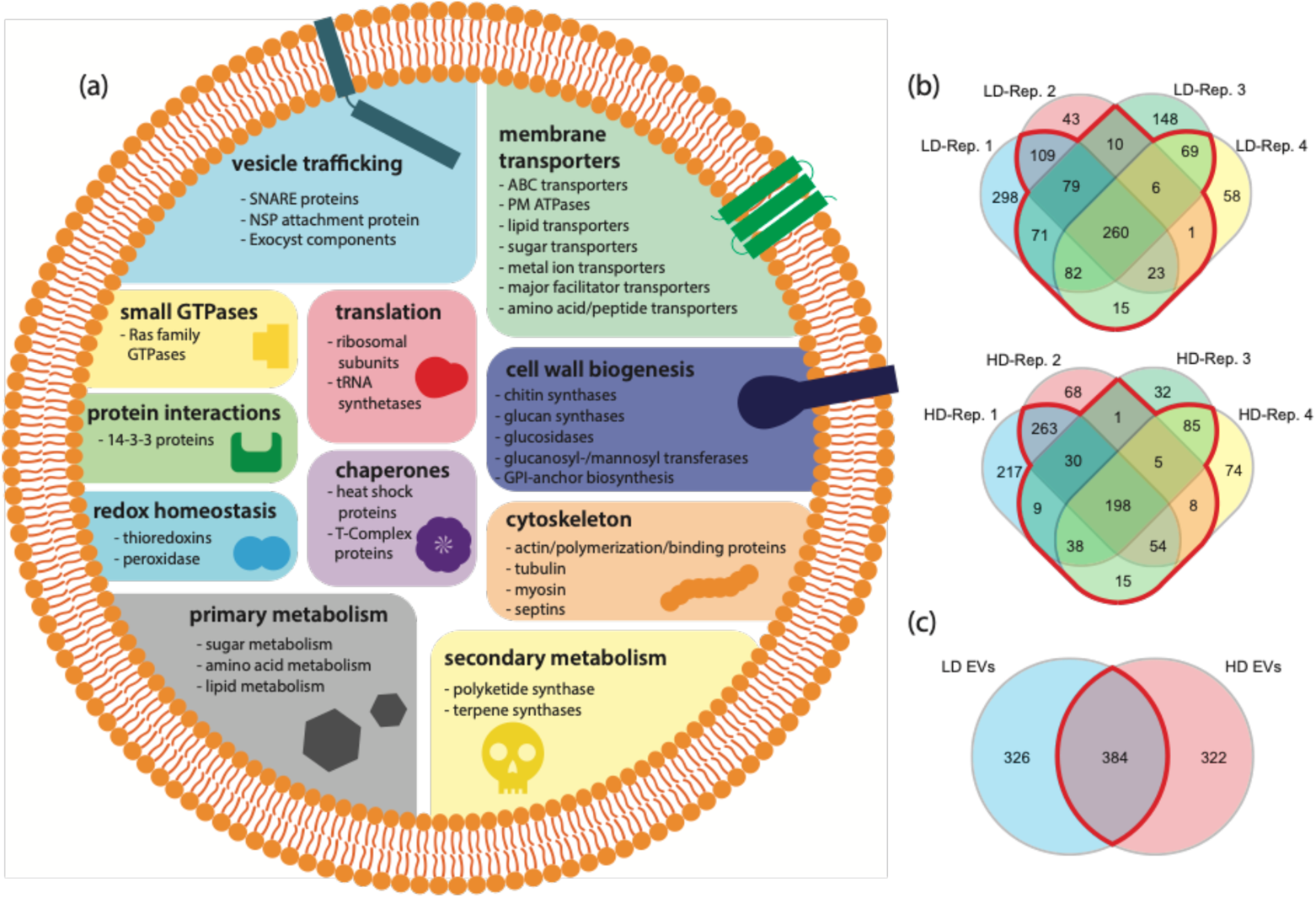
Overview of the *C. higginsianum* EV Proteome. (a) Overview of categories of proteins found in both low density (LD) and high density (HD) vesicles purified from *C. higginsianum* protoplasts. (b) Venn diagrams showing the overlap of proteins detected by mass spectrometry between independent replicates of purified LD and HD vesicles (proteins selected were present in 2 out of 4 replicates, q-value ≤ 0.01). The red outlined region contains protein shared by at least two replicates. (c) Venn diagram showing the overlap between the LD and HD vesicle proteomes (proteins selected were present in 2 out of 4 replicates, q-value ≤ 0.01). The red outlined region is shared. Approximately 46% of proteins in each population of vesicles are shared.

Mammalian EVs are associated with a common set of proteins that include transmembrane proteins (annexins and tetraspanins), vesicle trafficking proteins (ESCRT proteins and Rab GTPases), cytoskeletal proteins, heat-shock proteins, metabolic enzymes, integrins, 14-3-3 proteins and ribosomal subunits (Choi et al., 2015). Fungal EVs contain many similar proteins and are often enriched for proteins that function in the plasma membrane, cell wall biogenesis, pathogenesis, stress responses, transport, signaling and basic cellular metabolism (Bleackley, Dawson, et al., 2019). Despite the increasing number of studies on fungal EV proteins, scientists are still struggling to identify reliable protein biomarkers (Bleackley, Dawson, et al., 2019). A recent study attempting to define protein markers for *Candida albicans* EVs proposed 22 different proteins enriched in EVs compared to whole cell lysate. In addition to plasma membrane proteins, SNAREs, ER membrane proteins, cell surface proteins and GTPases, the list includes three Sur7 proteins, which are four transmembrane domain proteins localized in plasma membrane subdomains known as eisosomes in yeasts and filamentous fungi (Dawson et al., 2020). Interestingly, Sur7 proteins were also identified in EVs from the genus *Cryptococcus* and the wheat pathogen *Zymoseptoria tritici* (Hill & Solomon, 2020; Rizzo et al., 2021).

Between the two populations of vesicles from *C. higginsianum*, we identified several proteins commonly associated with EVs in both mammals and fungi. Based on GO term enrichment, the most enriched EV proteins in both fractions were membrane transporters (i.e. ABC and MFS transporters, plasma membrane proton ATPases), small GTPases, enzymes involved in amino acid and carbohydrate metabolism, as well as proteins involved in vesicle-mediated transport and translation (ribosomal subunits) (Figure 5a, Tables S1, S2). We also detected other classes of proteins involved in cytoskeleton organization, protein folding (heat-shock proteins), cell redox homeostasis and the synthesis of secondary metabolites (Figure 5a, Tables S1 to S3). While the majority of proteins were shared between the LD and HD fractions, the LD population was enriched for GO terms related to translation, amino acid metabolism, intercellular transport and actin filament polymerization (Table S4). The HD population was enriched for GO terms related to protein and lipid metabolism, protein folding and exocytosis, as well as membrane anchoring and docking (Table S4).

EVs are commonly associated with plasma membrane proteins (Bleackley, Dawson, et al., 2019). We found that approximately 19.7% and 35.4% of LD and HD vesicle proteins, respectively, had predicted transmembrane regions (Table S2). Membrane proteins associated with the vesicles included those involved in SNARE complexes (v-SNARES, t-SNARES and NSF attachment proteins), transport across membranes (ABC and MFS transporters, proton ATPases, sugar transporters, amino acid transporters, peptide transporters, lipid transporters) or recruitment of proteins to membranes (PH domain containing proteins, C2 domain proteins; Tables S1, S2). The vesicle proteomes also contained multiple GTPases (Ras, Rab, Ran and Rho GTPases) that function to regulate vesicle trafficking (Tables S1, S2). Interestingly, homologs of the *C. higginsianum* t-SNARE Sso2 (OBR10381) as well as the GTPases Ced-10 (OBR05828), RhoA (OBR05976), Rab-8A (OBR12345) Rab-11B (OBR03559), Arf3 (OBR07035) and Cell Division Control Protein 42 (OBR14052) were recently identified as EV markers in *C. albicans* (Dawson et al., 2020). These SNARES and GTPases were all present in the *C. higginsianum* LD and HD vesicle populations and could represent conserved fungal EV markers (Tables S1, S2).

Surprisingly, while we did detect Sur7-like proteins (OBR09999, OBR07542, OBR09056), they were not present at high enough peptide counts or in enough replicates to meet our criteria (Table S1). This suggests that, unlike *C. albicans* and members of the genus *Cryptococcus*, Sur7 proteins may not be enriched in *C. higginsianum* EVs. However, it should be noted that because our EV samples were pre-treated with trypsin, regions of Sur7 proteins on the surface of vesicles would have been degraded, making these proteins more difficult to detect by LC-MS/MS. For the same reason, it is possible that the calculated overall percentage of transmembrane proteins identified in both vesicle populations may have been underestimated.

A range of enzymes involved in cell wall synthesis and remodeling were detected in both populations of *C. higginsianum* vesicles. Fungal EVs are often associated with enzymes thought to function in cell wall synthesis and remodeling (Oliveira et al., 2010; Rizzo et al., 2020; Zhao et al., 2019). EVs from *S. cerevisiae* have the ability to promote cell wall remodeling in mutants with wall defects (Zhao et al., 2019), while EVs secreted from protoplasts of *A. fumigatus* are associated with fibrillar material and cell wall regeneration (Rizzo et al., 2020). EVs from *C. neoformans* were also observed passing through the cell wall, a process suggested to be made possible by enzyme-mediated cell wall remodeling (Wolf et al., 2014). Among LD and HD vesicle populations from *C. higginsianum*, we detected both chitin and glucan synthases, glucanosyl- and mannosyl-transferases, glucosidases and one GPI-anchored cell wall organizing protein (Tables S1, S2). The presence of these proteins suggests a role for *C. higginsianum* EVs in cell wall biology. Although the majority of these proteins were detected in both EV populations, the mannosyl transferases were enriched in the HD population, suggesting that different populations of vesicles could have distinct functions in cell wall remodeling.

We also detected proteins associated with the production of secondary metabolites (Table S3). Species of *Colletotrichum* as well as other pathogenic fungi produce a range of secondary metabolites, including polyketides, non-ribosomal peptides, alkaloids and terpenes. These metabolites often serve important roles in pathogenicity or virulence (Moraga et al., 2019). HD vesicles were associated with proteins involved in the production of higginsianin A to E. These metabolites are diterpenoid α-pyrones that have cytostatic and cytotoxic activities in mammalian cancer cells (Cimmino et al., 2016; Sangermano et al., 2019). In plants, higginsianin B inhibits the activation of jasmonate (JA) defense signalling by exogenous JAs as well as the wound-induced activation of this pathway (Dallery et al., 2020). Proteins involved in the higginsianin biosynthetic pathway present in the HD vesicle population included flavin-containing superfamily amine oxidase (OBR09790), terpene cyclase (OBR09784) and prenyltransferase (OBR09786; Table S3). Four other proteins in the pathway (geranylgeranyl pyrophosphate synthase (OBR09788), FAD-dependent epoxidase (OBR09783) and short-chain dehydrogenases (OBR09785 and OBR09787) were also detected but did not meet our criteria for inclusion in the final proteome (Table S1).

Proteins encoded by two additional secondary metabolite Biosynthetic Gene Clusters (BGC21 and BGC71) were also detected in LD and HD vesicles (Table S3). BGC21 comprises 12 genes encoding OBR10834 to OBR10845, including a polyketide synthase (PKS, OBR10843) and a terpene cyclase (OBR10835). This BGC is absent from other sequenced *Colletotrichum* spp. but homologs occur in diverse phylogenetically distant fungi (Figures S2a and S2b). No characterized chemical product has been described for any of the homologous BGCs, suggesting that BGC21 produces an unknown family of molecules (Table S5). BGC71 comprises 11 genes encoding OBR03047 to OBR03057 and is conserved in most *Colletotrichum* spp. and several Dothideomycetes, all of which are plant pathogens (Figures S3a and S3b). OBR03057 was detected in both LD and HD vesicles and together with OBR03051 and OBR03052, these three proteins resemble the lovastatin/monacolin BGC (Table S6). BGC71 may thus produce lovastatin-like molecules that could contribute to fungal pathogenicity.

Finally, we used SignalP-5.0 to predict the number of vesicle-associated proteins with N-terminal signal peptides (SPs). As an unconventional pathway for secretion, EVs usually have a lower percentage of proteins with SPs. For example, only 6.7% of EV proteins from the wheat pathogen *Zymoseptoria tritici* have a predicted SP (Hill & Solomon, 2020), while 12.5-22% of EV proteins from the cotton pathogen *Fusarium oxysporum f. sp. vasinfectum* had a predicted SP, depending on the medium in which the fungus was grown (Bleackley, Samuel, et al., 2019; Garcia-Ceron, Dawson, et al., 2021). Compared to these other filamentous plant pathogens, 4.9% of *C. higginsianum* LD vesicle proteins and 16.4% of HD vesicle proteins have a predicted signal peptide (Table S2).

### Development of Transgenic EV Marker Lines

Biomarkers are crucial tools in EV research; they can aid in the identification, authentication, tracking and purification of vesicles. In an effort to establish general protein biomarkers for *C. higginsianum* EVs, we selected protein candidates common to both the LD and HD populations that would serve as markers for vesicle membrane or luminal cargo. To mark the vesicle membrane, we selected the v-SNARE *Ch*Snc1 (OBR08408) and the t-SNARE *Ch*Sso2 (OBR10381), which are both integral membrane proteins and members of the Snc1/2, Sso1/2, sec9 SNARE complex. All members of this complex are present in the EV proteome. SNARE proteins are highly conserved across species, often maintaining the same function at different membrane trafficking steps (Bock et al., 2001; Gupta & Brent Heath, 2002; Kloepper et al., 2007). The Snc1/2, Sso1/2, Sec9 SNARE complex functions in the docking and fusion of exocytic vesicles with the plasma membrane (Aalto et al., 1993; Brennwald et al., 1994; Rossi et al., 1997). Orthologous SNARE proteins in other species of fungi have important roles in pathogenesis. For example, orthologs of *Ch*Sso2 mediate the secretion of pathogenesis-related degradation enzymes in *C. albicans* and the unconventional secretion of effectors in the phytopathogens *M. oryzae* and *V. dahliae* (Bernardo et al., 2014; Giraldo et al., 2013; Wang et al., 2018) and trichothecene phytotoxins in *Fusarium graminearum* (O’Mara et al., 2020). Moreover, orthologs of Snc1/2, Sso1/2 or Sec9 have been identified in the EV proteomes of *S. cerevisae*, *C. albicans*, *A. fumigatus* and *C. neoformans* (Dawson et al., 2020; Rizzo et al., 2020; Wolf et al., 2014; Zhao et al., 2019).

As a marker for EV luminal cargo, we selected the 14-3-3 protein *Ch*Bmh1 (OBR15609), one of two such proteins we detected in LD and HD vesicles. Members of the 14-3-3 family mediate protein-protein interactions and are involved in a myriad of cellular process, including cell division, vesicle trafficking, stress responses, signaling and apoptosis (Gelperin et al., 1995; Kumar, 2017; Roth et al., 1999; van Heusden & Steensma, 2006). 14-3-3 proteins are common in mammalian EVs (Choi et al., 2015), and have also been identified in the EV proteomes of several fungi, including *C. neoformans*, *S. cerevisae*, *F. oxysporum f. sp. vasinfectum*, *C. albicans*, *Z. tritici* and *A. fumigatus,* all of which possess at least one ortholog of *Ch*Bmh1 (Bleackley, Samuel, et al., 2019; Dawson et al., 2020; Hill & Solomon, 2020; Rizzo et al., 2020; Wolf et al., 2014; Zhao et al., 2019).

To test the association of our selected markers with *C. higginsianum* EVs and to validate our proteomic data, we generated transgenic lines expressing fluorescent-tagged marker proteins. *Ch*Snc1 and *Ch*Sso2 were expressed as N-terminal fusions with mScarlet and mNeonGreen, respectively, while *Ch*Bmh1 was expressed as a C-terminal fusion with mCherry. The mScarlet-*Ch*Snc1 construct was expressed from the native promoter, while the two others were placed under control of the *Ch*Actin promoter because their fluorescence signals were too weak or not detectable using their respective native promoters. The expression of each fluorescent fusion protein was verified in axenic mycelial cultures using confocal microscopy.

The mScarlet-*Ch*Snc1 fusion protein localized to the fungal plasma membrane and to small, mobile puncta that were concentrated near the hyphal apex (Figure 6a). Some hyphae also showed a small focal accumulation beneath the plasma membrane at the hyphal apex (Figure 6b), which likely corresponds to the Spitzenkörper (SPK), a cluster of microfilaments and vesicles that is pivotal for polar growth and apical secretion in fungi (Riquelme & Sanchez-Leon, 2014). In older, subapical regions, mScarlet-*Ch*Snc1 strongly labelled septa but the PM labelled less strongly than in apical regions and vacuoles were labelled weakly (Figure 6c). A similar localization of Snc1 proteins was also observed in hyphae of *Fusarium graminearum* (Zheng et al., 2018), *Magnaporthe oryzae* (Giraldo et al., 2013) and *Aspergillus oryzae* (Hayakawa et al., 2011; Kuratsu et al., 2007). The mNeonGreen-*Ch*Sso2 fusion protein predominantly labeled septa and the plasma membrane in subapical regions, with fluorescence intensity progressively decreasing towards the hyphal tip (Figures 6d and e). A similar subapical PM localization of Sso2 was also reported in *A. oryzae* (Kuratsu et al., 2007). Finally, *Ch*Bmh1-mCherry appeared uniformly distributed through the cytoplasm of hyphae but was excluded from vacuoles. In some cases, a small apical region, probably corresponding to the SPK, was labelled more strongly than the rest of the cytoplasm (Figures 6f and 6g).

**FIGURE 6:**
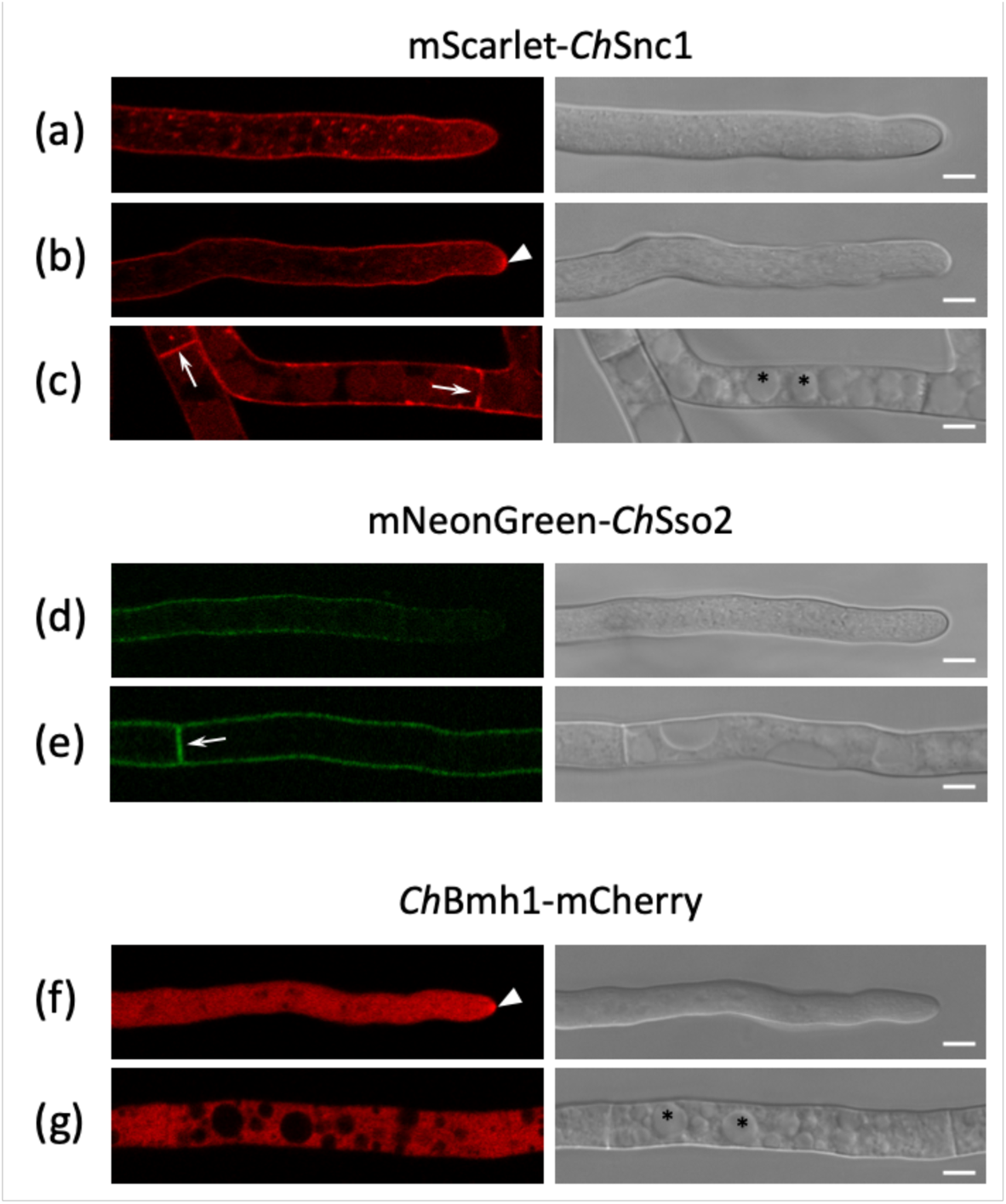
Localization of EV marker proteins in vegetative hyphae on Mathur’s agar. Confocal laser scanning microscope images showing fluorescence and DIC channels. mScarlet-*Ch*Snc1 localized to the fungal plasma membrane (PM) and small, mobile punctae concentrated near the hyphal apex (a), as well as the Spitzenkörper (SPK, arrowhead) (b). In older, subapical regions, mScarlet-*Ch*Snc1 strongly labelled septa (arrows) but the PM labelled less strongly than in apical regions. Vacuoles (asterisks) were labelled weakly (c). mNeonGreen-*Ch*Sso2 labelled septa (arrow) and the plasma membrane in subapical regions (e), with fluorescence intensity progressively decreasing towards the hyphal tip (d). *Ch*Bmh1-mCherry was distributed throughout the hyphal cytoplasm but was excluded from vacuoles (f, g, asterisks) and in some hyphae the SPK (arrowhead) was strongly labelled (f). Scale bars = 5 µm.

### Fluorescent EV Marker Proteins are Protected Inside EVs

To validate the association of our selected marker proteins with *C. higginsianum* EVs, we grew mycelial cultures of each transgenic line alongside wild-type, non-transgenic *C. higginsianum*. Protoplasts were generated from each culture, and crude EV samples were isolated from the supernatant. Equal amounts of protein from samples of whole-cell lysate and EVs were probed for fluorescent-tagged proteins using an immunoblot. We detected mScarlet-*Ch*Snc1, *Ch*Bmh1-mCherry and mNeonGreen-*Ch*Sso2 by immunoblot in lysate and vesicle samples from transgenic fungi but not in samples from the non-transgenic fungus (Figure 7). Both mScarlet-*Ch*Snc1 and mNeonGreen-*Ch*Sso2 appeared to be enriched in EV samples compared to whole-cell lysate. It makes sense that both SNARE proteins would be enriched, because they interact in the same complex. However, the signal for mNeonGreen-*Ch*Sso2 was much weaker than that of mScarlet-*Ch*Snc1. This lower level of expression could be a result of the mNeonGreen tag, which was not codon optimized for this fungus. We also probed vesicle samples for native Superoxide Dismutase 2 (SOD2), a mitochondrial protein that was recently identified as a potential negative EV marker (Dawson et al., 2020). While we detected SOD2 in samples of mycelial lysate, we were unable to detect it in vesicle samples from transgenic or non-transgenic fungi (Figure 7). These results show that vesicle samples are enriched for our selected EV markers but not a marker for contamination, indicating that these vesicles are unlikely to be the product of cell lysis.

**FIGURE 7:**
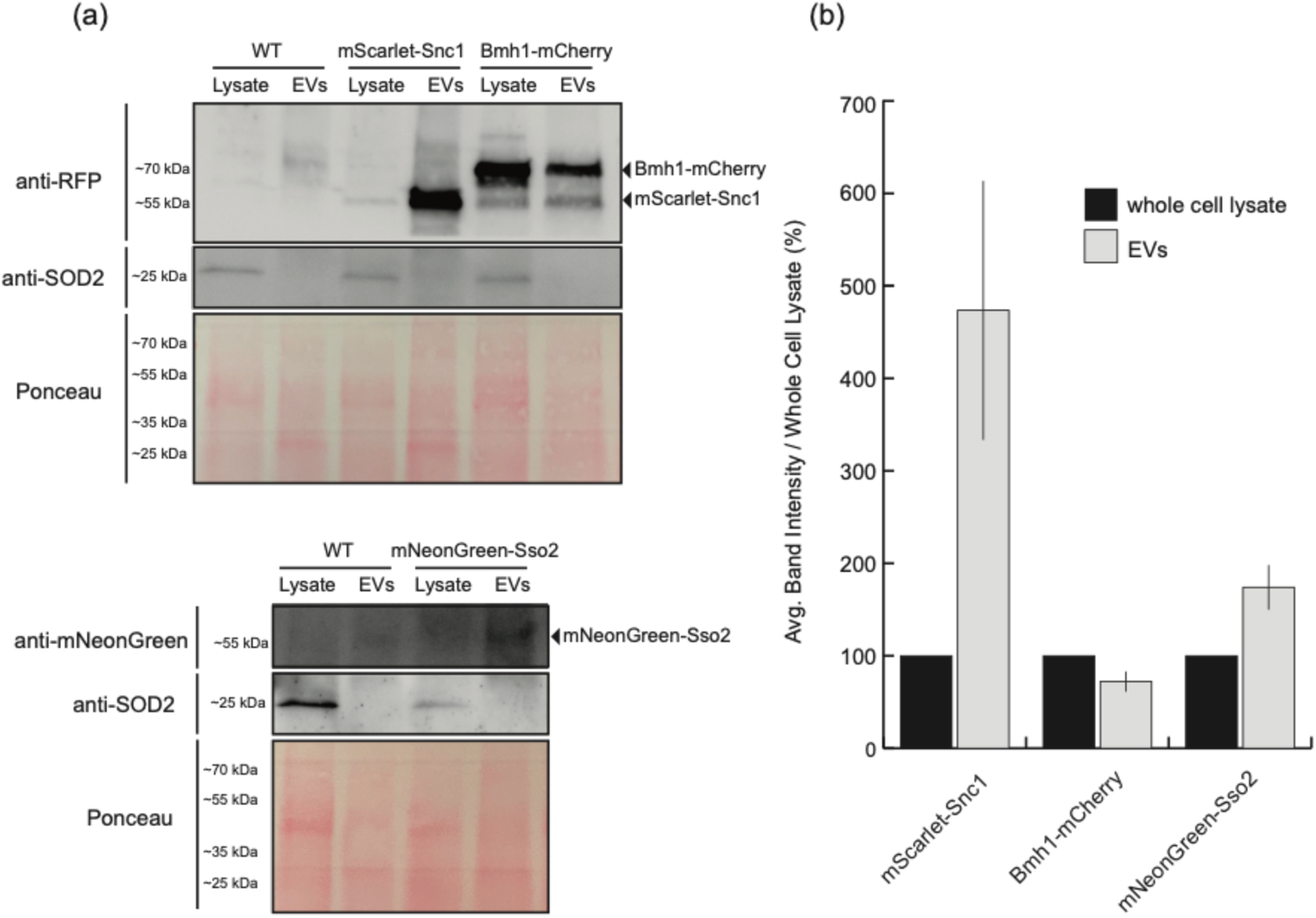
Detection of transgenic markers in EV samples. (a) Crude vesicle pellets from transgenic marker lines and nontransgenic wild-type fungi were isolated and probed for the fluorescent-tagged proteins RFP or mNeonGreen by immunoblot. EV markers were detected in samples of protoplast lysate as well as crude vesicle samples, while the contamination marker SOD2 was only detected in the sample of mycelial lysate. (b) EV marker band intensities were quantified and expressed as a percentage of the band intensity for the whole cell lysate. Bars represent the average of two independent replicates.

Finally, we wanted to confirm that our fluorescent marker proteins were present inside fungal EVs rather than merely co-pelleting with the vesicles. To do this, we subjected vesicle samples to a protease protection assay. Vesicle samples were treated with either buffer, trypsin or detergent followed by trypsin. Proteins protected inside of vesicles are expected to remain intact after trypsin treatment unless detergent has been added to solubilize lipid membranes. Again, our marker proteins were detected in the lysate of transgenic mycelia and in crude vesicle samples. Importantly, the proteins were still detected in vesicle samples after treatment with trypsin but could not be detected when the samples were pre-treated with detergent followed by trypsin (Figure 8). These results indicate that the marker fusion proteins are packaged inside of fungal EVs, where they are protected from degradation. This helps validate our proteomic data and suggests that *Ch*Snc1, *Ch*Sso2 and *Ch*Bmh1 can all function as markers for *C. higginsianum* EVs. However, mNeonGreen-*Ch*Sso2 may require additional optimization to be reliably detected in fungal mycelia and EVs.

**FIGURE 8:**
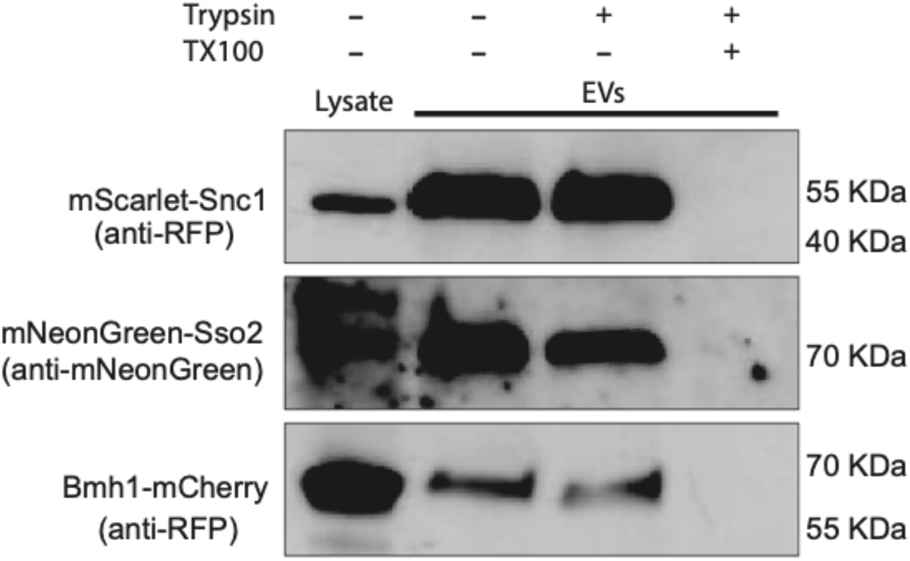
Transgenic EV markers are protected inside of lipid vesicles. A sample of crude vesicles from each EV marker line was split into three and either left untreated, treated with trypsin or treated with detergent followed by trypsin. EV markers were then detected by immunoblot. Detection of the fusion protein in the presence of trypsin but not in the presence of detergent plus trypsin indicates the protein is protected within a lipid compartment. All experiments were repeated twice.

### Localization of EV Markers During Infection

One of the reasons for developing the above EV marker lines was to enable tracking of EVs during infection of plants. To this end, we infected *Arabidopsis* leaves with our transgenic EV marker lines and observed the localization of each marker at different stages of development. Early in infection, the *C. higginsianum* spore germinates and produces a nascent, pre-penetration appressorium devoid of melanin. Over time, the appressorium matures and its cell wall becomes melanized. The base of this specialized structure contains a penetration pore from which a needle-like penetration peg emerges to pierce the cuticle and plant cell wall. After the fungus successfully penetrates the plant cell wall, it produces a bulbous, multi-lobed biotrophic hypha. Eventually, the fungus transitions to a necrotrophic stage, producing long, thin necrotrophic hyphae capable of invading and overwhelming neighboring cells (O’Connell et al., 2004). We examined the localization of markers in spores, appressoria, biotrophic hyphae and necrotrophic hyphae.

In spores expressing mScarlet-*Ch*Snc1, we observed numerous punctate bodies that were possibly small vacuoles (Figure S4). In appressoria, mScarlet-*Ch*Snc1 signals were also observed in small fluorescent puncta arranged around the cell periphery (Figures 9a and 9b), and in some young appressoria the plasma membrane was also weakly labelled (Figure S4). Within young biotrophic hyphae, mScarlet-*Ch*Snc1 was uniformly distributed in the plasma membrane and in small puncta (Figures 9c and 9d), whereas in more mature biotrophic hyphae the plasma membrane labelling was more concentrated at hyphal tips (Figures 9e and 9f). After the transition to necrotrophy, mScarlet-*Ch*Snc1 was similarly localized to the apical plasma membrane of necrotrophic hyphae, and numerous vacuoles of varying sizes were also intensely labelled (Figures 9g to 9j). The fluorescent signal for mNeonGreen-*Ch*Sso2 was relatively weak, even when expressed from the actin promoter, and no signals were detectable in either spores or appressoria. However, in biotrophic hyphae, mNeonGreen-*Ch*Sso2 was localized to the plasma membrane as well as to septa (Figures 10c and 10d). Labelling of the plasma membrane with mNeonGreen-*Ch*Sso2 appeared weaker in necrotrophic hyphae compared to biotrophic hyphae (Figures 10a and 10b). Similar to mScarlet-*Ch*Snc1, *Ch*Bmh1-mCherry localized to small punctate bodies in spores (Figure S4), appressoria (Figures 11a and 11b), and young biotrophic hyphae (Figures 11c to 11f). However, the fusion protein showed a strikingly different localization in more mature biotrophic hyphae (Figures 11g and 11h) and necrotrophic hyphae (Figures 11i and 11j), where *Ch*Bmh1-mCherry was instead distributed throughout the cytoplasm, but excluded from vacuoles. At no point during the infections were we able to observe potential extracellular signals for our EV markers, neither in the plant apoplast nor inside the plant cytoplasm. Capturing such signals may require more sensitive imaging techniques or sampling at other timepoints. However, mScarlet-*Ch*Snc1 was consistently concentrated at hyphal apices, which are regions of intense polarized secretion associated with the SPK (Riquelme & Sanchez-Leon, 2014). Whether the localization of mScarlet-*Ch*Snc1 to this region implies EV secretion occurs at hyphal tips or only the presence of abundant secretory vesicles remains to be determined.

**FIGURE 9:**
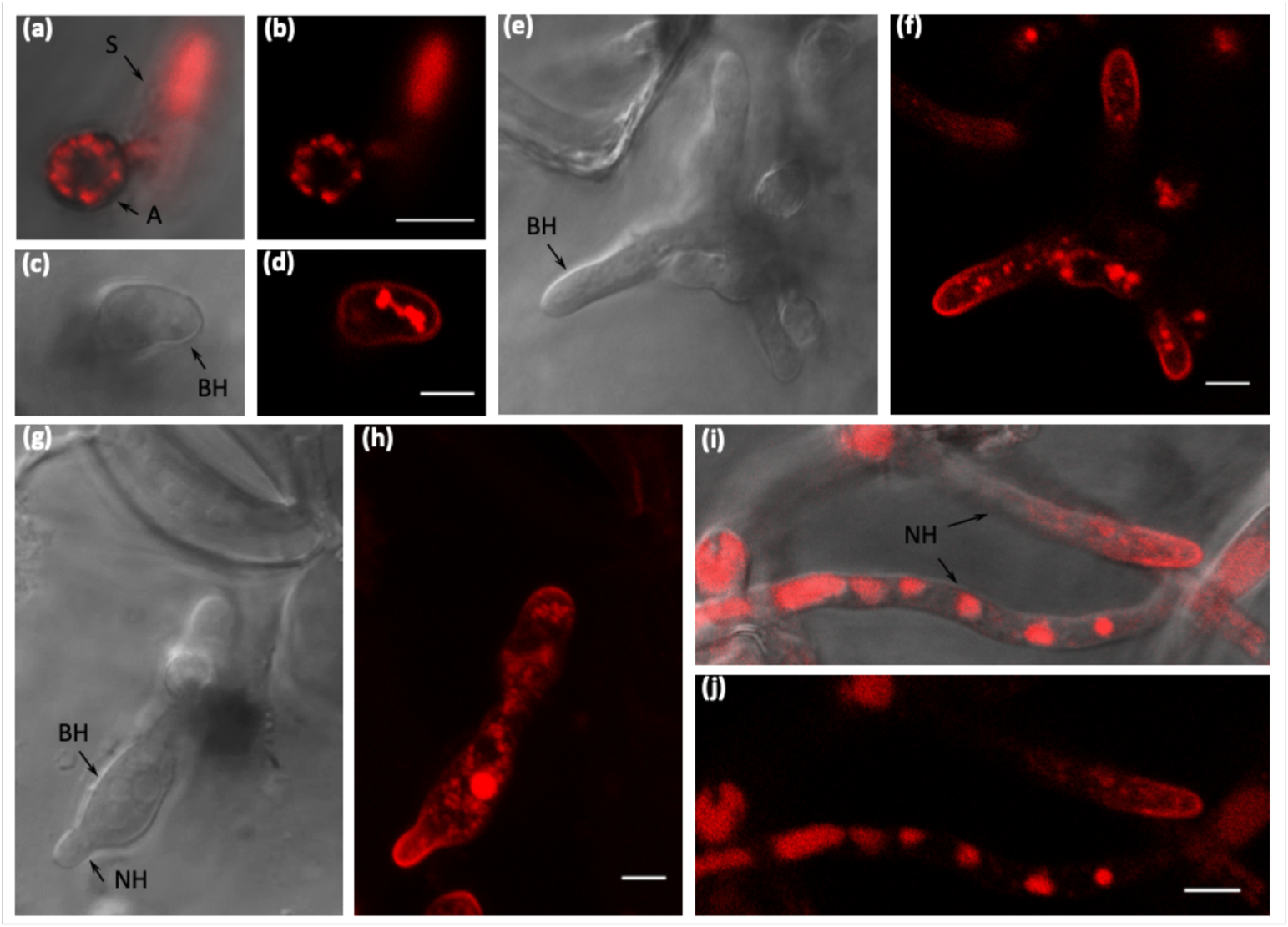
Localization of mScarlet-ChSnc1 in *C. higginsianum* infecting Arabidopsis. Fungal transformants expressing mScarlet-*Ch*Snc1 were inoculated on Arabidopsis cotyledons and observed with confocal laser scanning microscopy. (a, b) mScarlet-*Ch*Snc1 labelled vacuoles in spores and small punctae in appressoria. (c-f) In young biotrophic hyphae, mScarlet-*Ch*Snc1 localized to the plasma membrane (PM) and small punctae. (g-h) At the transition to necrotrophy, mScarlet-*Ch*Snc1 labelled the PMs of both biotrophic and necrotrophic hyphae. (I, J) In necrotrophic hyphae, mScarlet-*Ch*Snc1 localized to the PM at hyphal tips and strongly labelled vacuoles. (b, d, f, h, j) fluorescence channel; (c, e, g) DIC channel; (a, i) Overlays of DIC and fluorescence channels. Scale bars = 5 µm. S, spore; A, appressorium; BH, biotrophic hypha; NH, necrotrophic hypha.

**FIGURE 10:**
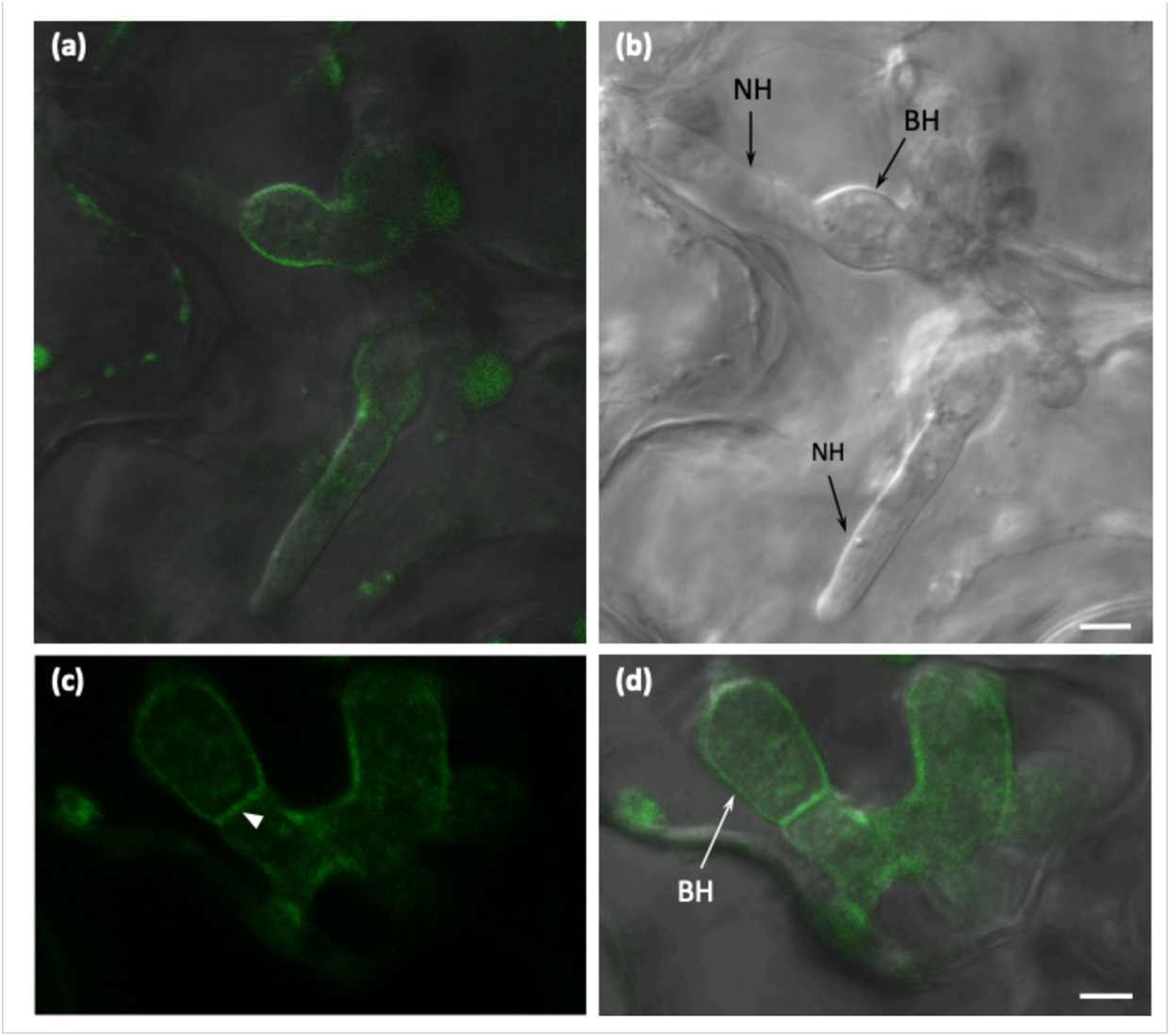
Localization of mNeonGreen-*Ch*Sso2 in *C. higginsianum* infecting Arabidopsis. Fungal transformants expressing mNeonGreen-*Ch*Sso2 were inoculated on Arabidopsis cotyledons and observed with confocal laser scanning microscopy. In biotrophic hyphae (BH), mNeonGreen-*Ch*Sso2 localized to the plasma membrane (PM) as well as to septa (arrowhead). PM labelling appeared weaker in necrotrophic hyphae (NH) than in biotrophic hyphae. (a, d) Overlays of DIC and fluorescence channels; (b) DIC channel; (c) fluorescence channel. Scale bars = 5 µm.

**FIGURE 11:**
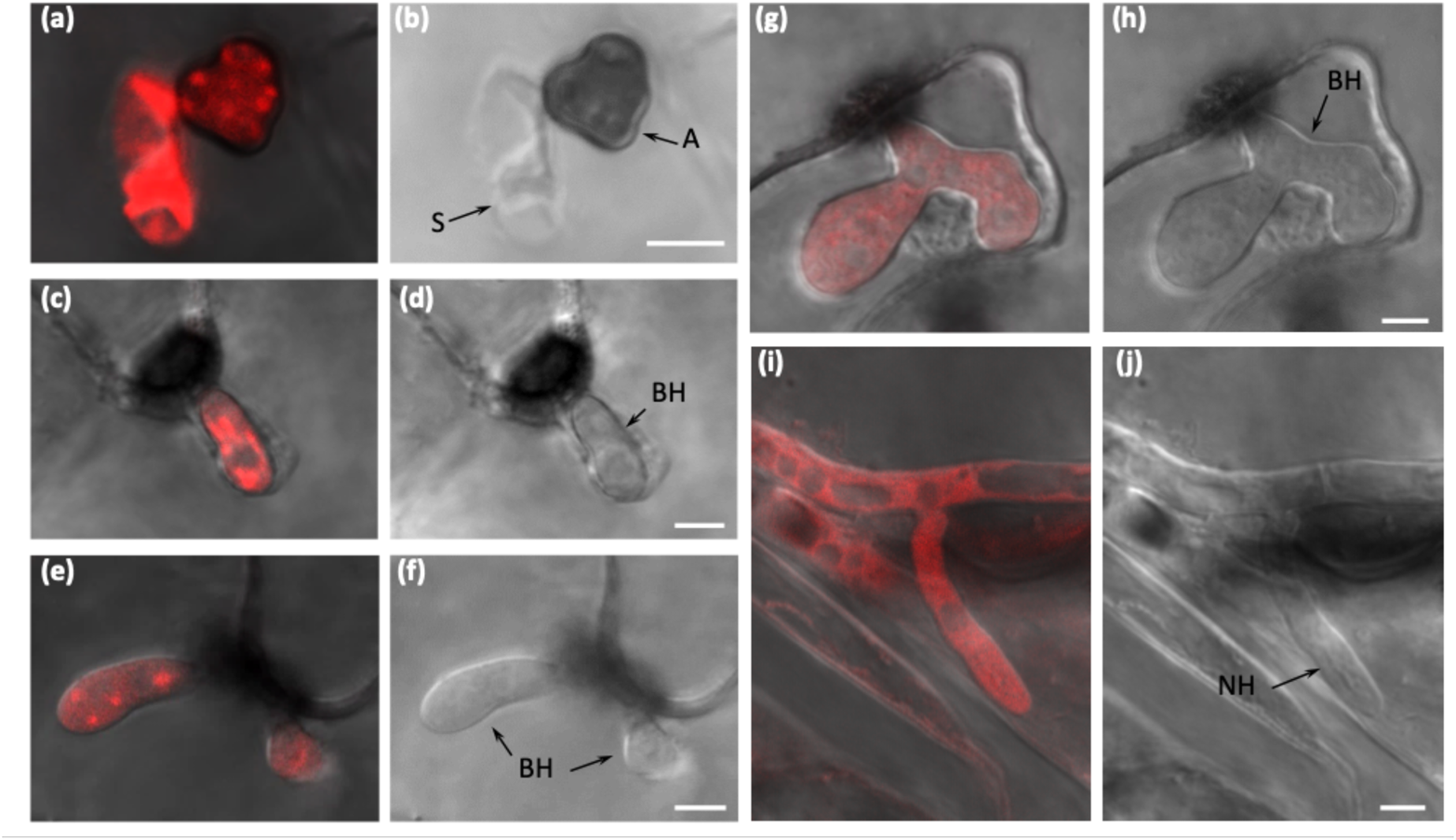
Localization of ChBmh1-mCherry in *C. higginsianum* infecting Arabidopsis. Fungal transformants expressing ChBmh1-mCherry were inoculated on Arabidopsis cotyledons and observed with confocal laser scanning microscopy. (a, b) ChBmh1-mCherry labelled vacuoles in spore and punctae in appressorium. (c-f) ChBmh1-mCherry localized to small punctae in young biotrophic hyphae. (g-j) In older biotrophic and necrotrophic hyphae, fluorescent labelling was distributed through the cytoplasm and excluded from vacuoles. (a, c, e, g, i) Overlays of DIC and fluorescence channels; b, d, f, h, j) DIC channel. Scale bars = 5 µm. S, spore; A, appressorium; BH, biotrophic hypha; NH, necrotrophic hypha.

## Discussion

Fungal plant pathogens rely on large secretomes to successfully infect plants (Krijger et al., 2014). While much of the fungal secretome is released through conventional mechanisms, a large percentage of proteins secreted by fungal phytopathogens can exit the cell through unconventional pathways (Giraldo et al., 2013; Kim et al., 2013; Nogueira-Lopez et al., 2018; Wang et al., 2018). Approximately, 54% of apoplastic proteins secreted by the maize root pathogen *Trichoderma virens* lack a signal peptide (Nogueira-Lopez et al., 2018). The same is true for 48% of the total secretome of *M. oryzae* (Kim et al., 2013). As a form of unconventional secretion, EVs represent an attractive mechanism for the release of important fungal virulence factors. EVs are frequently observed by TEM at the interface between plant and fungal cells (Ivanov et al., 2019; Micali et al., 2011; Roth et al., 2019). Recent studies also suggest that EVs from fungal phytopathogens carry protein effectors and phytotoxic compounds (Bleackley, Samuel, et al., 2019; Garcia-Ceron, Dawson, et al., 2021; Garcia-Ceron, Lowe, et al., 2021).

To better understand the composition and function of EVs from plant fungal pathogens, we developed methods for isolating EVs from the hemibiotrophic pathogen, *C. higginsianum*. Our TEM analysis revealed EVs in the paramural space of biotrophic hyphae growing *in planta*, but attempts to isolate these structures from the supernatant of liquid cultured mycelia resulted in few if any particles. In order to isolate large numbers of vesicles from the supernatant, we found it necessary to first degrade the fungal cell wall with a mixture of β-glucanases. This suggests that *C. higginsianum* EVs do not pass efficiently through the fungal cell wall. Alternatively, these EVs may not cross the cell wall at all. They could instead function as a mechanism for bulk export to the paramural space, where they subsequently rupture, releasing their contents.

EVs were previously isolated from protoplasts of *Aspergillus fumigatus* by Rizzo et al. (2020). In that study, EVs were actively released from protoplasts, especially under conditions that favored cell wall regeneration. Earlier studies have also observed the release of vesicles from protoplasts of *Aspergillus nidulans* and *Candida tropicans* (Gibson & Peberdy, 1972; Osumi, 1998). It is unknown whether the EVs collected from *C. higginsianum* protoplasts were secreted in response to the loss of the cell wall or if they were merely released from the space between the plasma membrane and cell wall after the latter’s removal. We are, however, confident, that the isolated vesicles represent legitimate EVs that are not artifacts produced from burst hyphal tips or damaged protoplasts, as indicated by the lack of particles in the supernatant of intact mycelia, the low level of cell death in samples of protoplasts and the absence of the contamination marker SOD2 in EV pellets.

Density gradient purification of *C. higginsianum* EV samples yielded two separate populations of vesicles, a low-density (LD) population at ~1.078 g/ml and a high-density (HD) population at ~1.115-1.173 g/ml. These vesicles were approximately 100 nm in diameter and, using TEM, had a similar appearance to EVs isolated from other fungal phytopathogens (Bleackley, Samuel, et al., 2019; Hill & Solomon, 2020; Kwon et al., 2021). In examining the protein content of these vesicles, we treated the vesicles with trypsin and then purified them on a density gradient, removing extra-vesicular proteins through both enzymatic digestion and density purification.

Our proteomic analysis of *C. higginsianum* EVs identified a diverse set of proteins for both vesicle populations, including membrane transporters, small GTPases, chaperones and ribosomal subunits, as well as proteins involved in primary/secondary metabolism, cell redox homeostasis, vesicle-mediated transport and cytoskeletal organization. Over 50% of the proteins detected in each vesicle population were held in common. However, the LD population was more enriched for cytoplasmic elements while the HD vesicles were enriched for membrane-associated proteins and contained a higher percentage of proteins with predicted trans-membrane domains. This difference could signify separate origins for each population.

Among the membrane-associated proteins identified, we detected SNAREs and GTPases previously proposed as fungal EVs markers for *C. albicans* (Dawson et al., 2020). The proteomes also contained multiple proteins associated with cell wall biogenesis, suggesting that, similar to other species of fungi, *C. higginsianum* EVs may function in the formation or maintenance of this structure (Oliveira et al., 2010; Rizzo et al., 2020; Zhao et al., 2019). Interestingly, we also identified proteins belonging to previously identified secondary metabolite biosynthetic gene clusters (BGCs), including proteins involved in the synthesis of higginsianins and two other as-of-yet unidentified compounds. Polyketide synthases have been found in the EVs of *Alternaria infectora* and *Fusarium oxysporum f. sp. vasinfectum*; EVs of the latter were also associated with an unidentified purple-colored phytotoxic metabolite (Bleackley, Samuel, et al., 2019; Silva et al., 2014). Proteins involved in toxin synthesis were also identified in EVs from *Fusarium graminearum* (Garcia-Ceron, Lowe, et al., 2021). Combined, these findings suggest that EVs may represent a common mechanism among fungal phytopathogens for the secretion, or even extracellular synthesis, of biologically active metabolites.

From our list of EV proteins, we selected two SNARE proteins (*Ch*Snc1 and *Ch*Sso2) and one 14-3-3 protein (*Ch*Bmh1) to function as markers for the EV membrane and lumen, respectively. Orthologs of these proteins have been identified in EVs from multiple species of fungi (Bleackley, Samuel, et al., 2019; Dawson et al., 2020; Hill & Solomon, 2020; Rizzo et al., 2020; Wolf et al., 2014; Zhao et al., 2019). All three markers were detected in EV samples by immunoblot, with *Ch*Snc1 and *Ch*Sso2 more enriched in EVs compared to total cell lysate. Moreover, a protease protection assay confirmed that all three markers were protected from digestion inside membranous compartments. In transgenic fungi grown on *Arabidopsis* leaves, we observed that *Ch*Snc1 was consistently localized to the apices of hyphae, known regions of secretion, while *Ch*Bmh1 had a more diffusive, cytoplasmic signal. Both markers localized to puncta inside appressoria. While fluorescent-tagged EV marker lines are available in yeasts (Dawson et al., 2020), to our knowledge, these are the first fluorescent EV marker lines for a filamentous plant pathogen. Though we could not detect extracellular signals with our current techniques, we hope these lines will prove valuable in the future for verifying EVs and tracking their secretion during infection.

Fungal EVs are a rapidly growing field of study. While the majority of research has focused on human pathogens, there is a dawning appreciation for the role of EVs among fungal phytopathogens. We present here, for the first time, an analysis of EVs produced by the hemibiotrophic plant pathogen, *C. higginsianum*. We found it was necessary to generate protoplasts to isolate EVs from the supernatant of liquid cultures. This suggests that *C. higginsianum* EVs do not pass efficiently through the fungal cell wall, if at all. This is most likely a problem common to all fungal EV studies. The majority of published fungal EV proteomes contain fewer than 100 proteins and sometimes pool EV samples from multiple independent replicates to achieve higher numbers (Bleackley, Dawson, et al., 2019; Bleackley, Samuel, et al., 2019). While the number of proteins detected depends on several factors ranging from the growth conditions to the type of equipment and software used for LC-MS/MS, the fungal cell wall undoubtedly represents an additional factor. It is already clear that external capsular structures are a barrier for the release of EVs from *C. neoformans* (Rodrigues et al., 2007). As previously suggested, protoplasting may therefore represent a more efficient way to determine the proteomes of fungal EVs (Rizzo et al., 2020).

Our research suggests that *C. higginsianum* secretes EVs that could function in cell wall biogenesis and the secretion of secondary metabolites. The proteins *Ch*Snc1 and *Ch*Bmh1 function as general markers for these vesicles and could be used in the future to verify and potentially track EVs produced by this fungus. These proteins are also attractive targets for genetic analysis, in order to determine how they impact the formation and contents of fungal EVs. Most excitingly, *C. higginsianum* is a pathogen of the model plant *Arabidopsis thaliana*, the EVs of which are currently the best-defined among plant species (Baldrich et al., 2019; Karimi et al., 2021; Rutter & Innes, 2017). Future research will need to determine if *C. higginsianum* secretes larger numbers of EVs past the fungal cell wall during infection. The tools developed here could be used to verify the presence of fungal EVs or EV-delivered cargo in the plant extracellular spaces. This could in turn lead to an exploration of how fungal and plant EV contents change during different phases of infection. A solid understanding of EVs secreted by both pathogen and host paves the way for future studies examining EV inter-kingdom communication.

## Acknowledgements

We thank the Indiana University Bloomington Light Microscopy Imaging Center and Electron Microscopy center, as well as the INRAE Versailles Plant Cell Imaging Platform for access to light and electron microscopes. We also thank Dr. Jonathan Trinidad and the Laboratory for Mass Spectrometry at Indiana University for the proteomic analysis and help describing the methods involved. We are also grateful to Adeline Simon and the BIOGER bioinformatics platform for advice on gene ontology analysis. Richard O’Connell is supported by funding from the Agence Nationale de la Recherche (grant ANR-17-CAPS-0004-01). The INRAE BIOGER unit benefits from the support of the Saclay Plant Sciences-SPS (ANR-17-EUR-0007). Roger Innes is supported by funding from the United States National Science Foundation (grant numbers IOS-1645745 and IOS-1842685).

## Supplemental Figures

**FIGURE S1:**
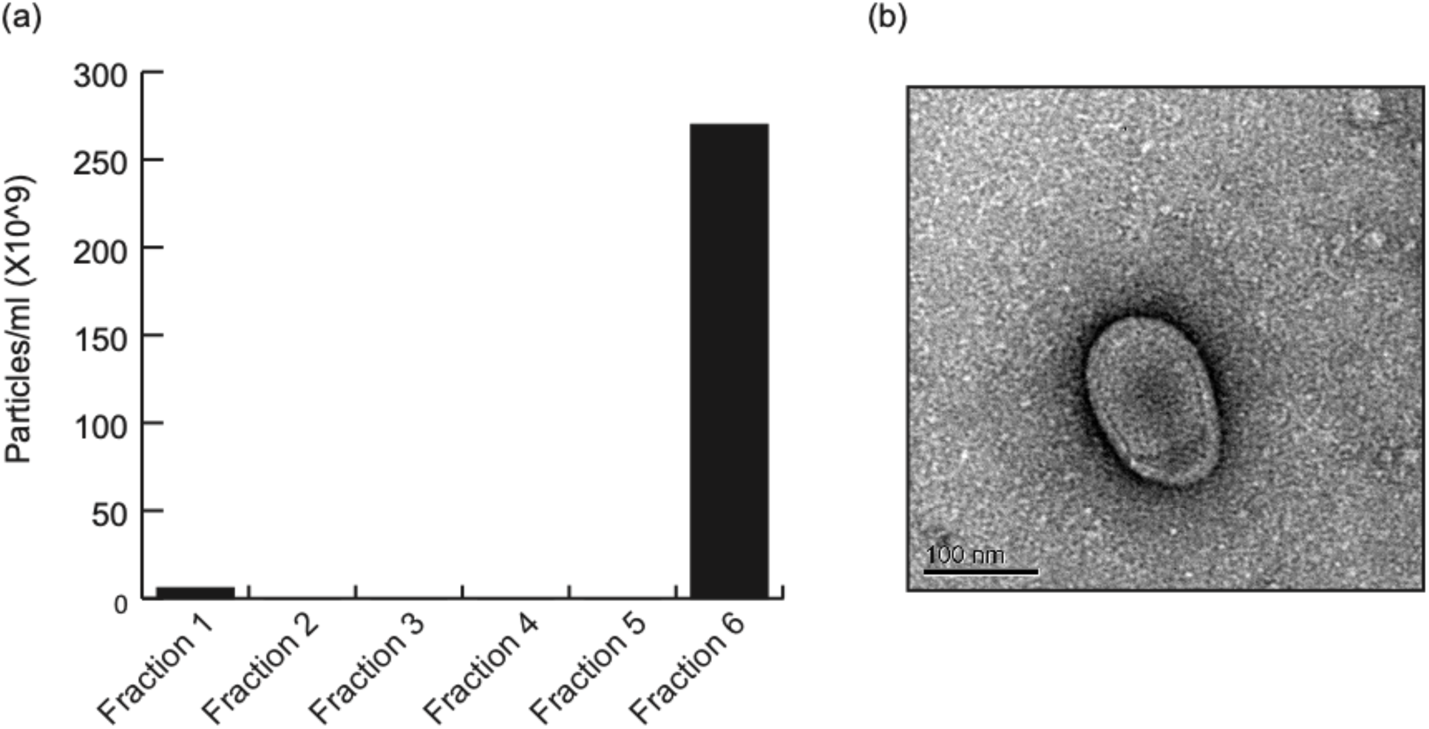
Preliminary attempts at isolating EVs from the supernatant of *C. higginsianum* mycelia. Supernatant from a mycelial liquid culture was processed for EVs. Crude vesicle pellets were bottom-loaded into a discontinuous Optiprep gradient consisting of 5, 10, 20 and 40% layers. After centrifugation at 100K × g for 17 hrs, the 5% layer was discarded and the next six fractions of 1 ml each were collected and processed with further ultracentrifugation. (a) Nanoparticle tracking (NTA) data showing the average concentration of particles in each of the collected Optiprep fractions. (b) TEM image of an EV-like particle found in fraction 6.

**FIGURE S2:**
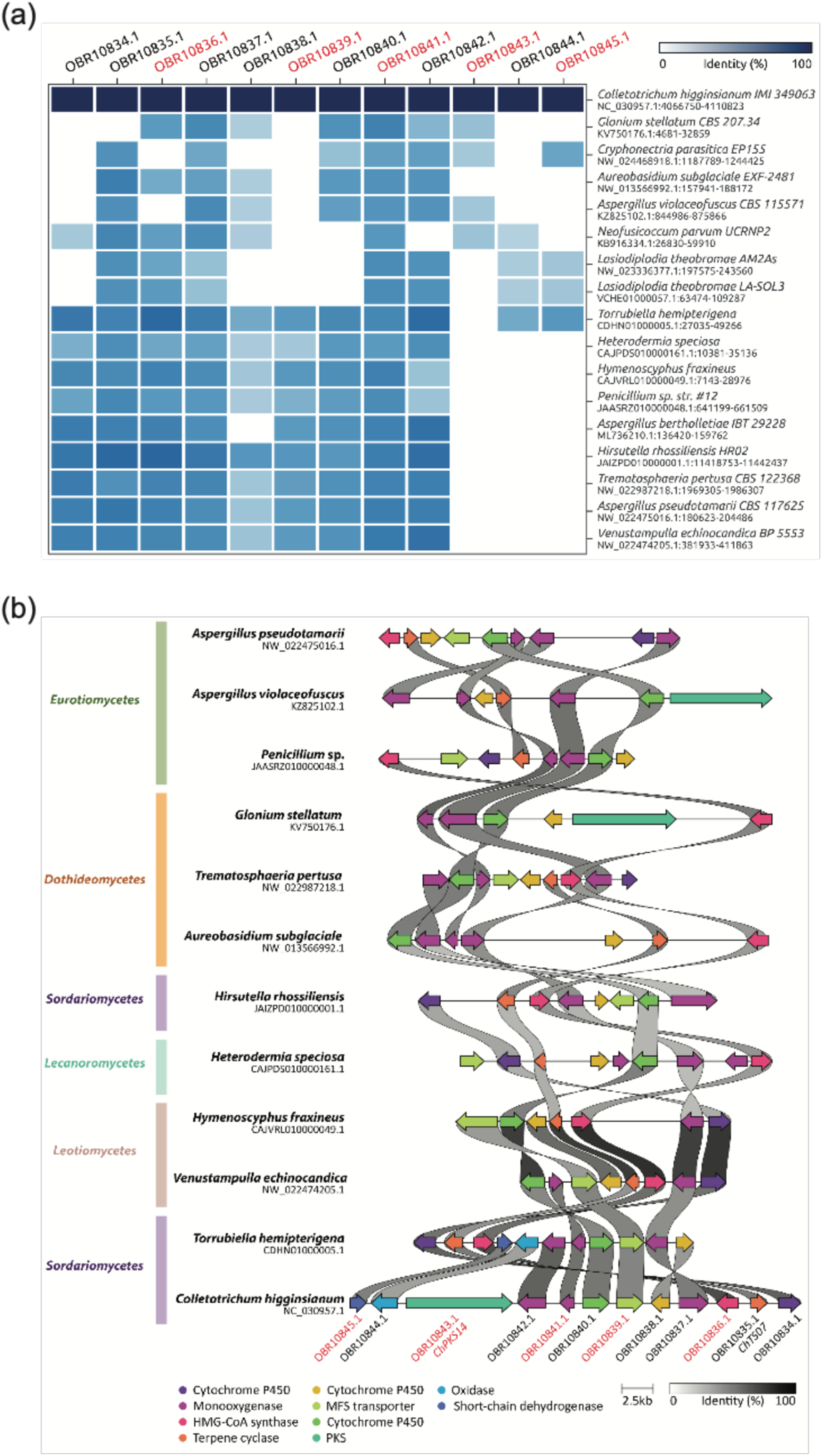
Homologous biosynthetic gene clusters of BGC21 from *C. higginsianum*. (a) Heatmap of the hits detected using cblaster v1.3.9 (Gilchrist et al., 2021a) with default parameters and (b) rearrangement of BGCs depicted using clinker v0.0.21 (Gilchrist et al., 2021b). IDs in red correspond to proteins detected in at least two biological replicates of EVs preparations.

**FIGURE S3:**
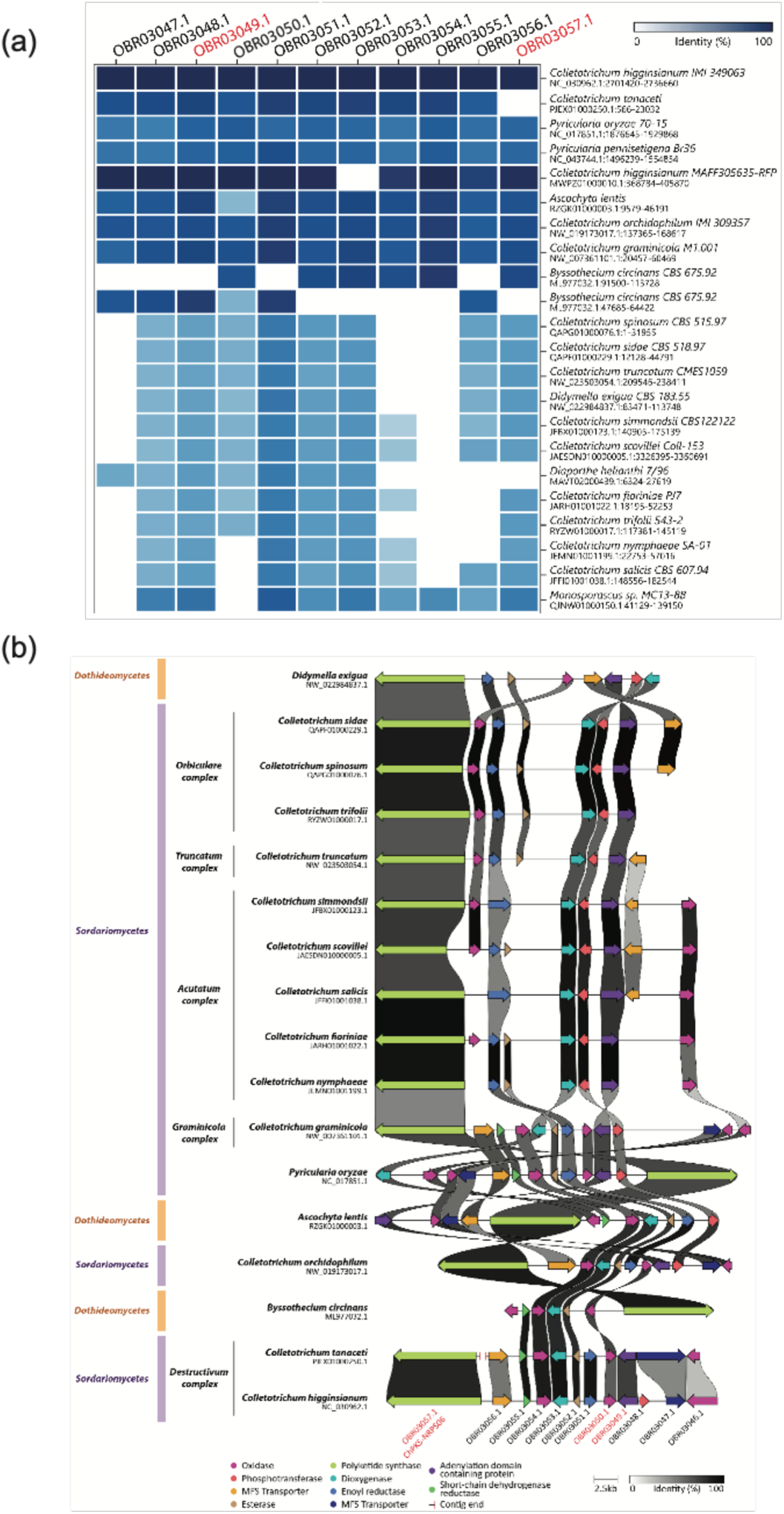
Homologous biosynthetic gene clusters of BGC71 from *C. higginsianum*. (a) Heatmap of the hits were detected using cblaster v1.3.9 (Gilchrist et al., 2021a) with default parameters and (b) rearrangement of BGCs depicted using clinker v0.0.21 (Gilchrist et al., 2021b). IDs in red correspond to proteins detected in at least two biological replicates of EVs preparations. Species complexes were indicated where appropriate for *Colletotrichum* spp.

**FIGURE S4:**
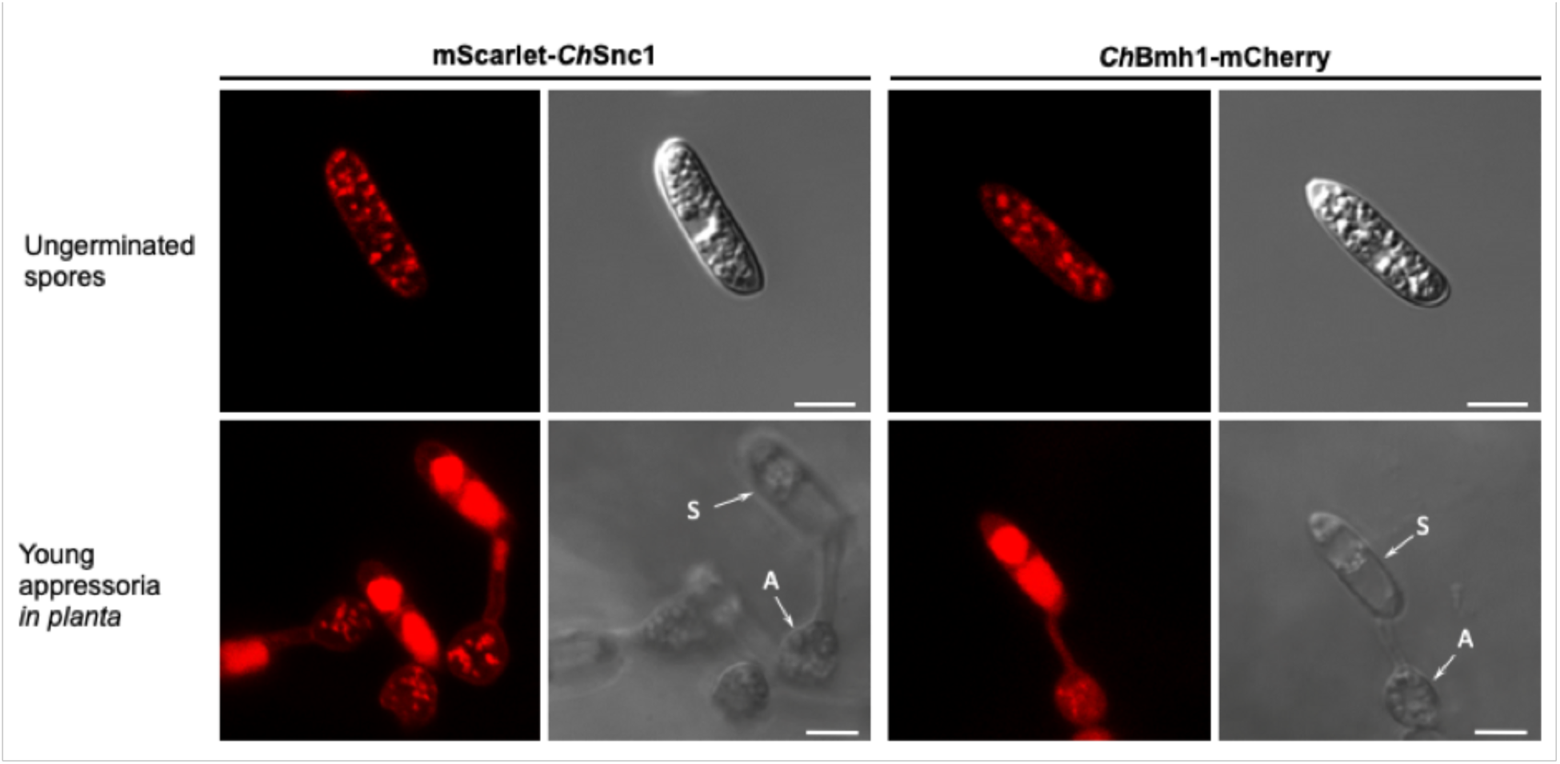
Localization of EV marker proteins in spores and young appressoria *in planta*. Confocal laser scanning microscope images showing ungerminated spores of *C. higginsianum* and germinated spores forming young appressoria (arrowheads) on the surface of Arabidopsis cotyledons (7 h post-inoculation), expressing either mScarlet-*Ch*Snc1 or *Ch*Bmh1-mCherry. In ungerminated spores, both fluorescent markers localize to abundant punctate bodies in the cytoplasm. Following germination on the plant surface, both markers label large vacuoles inside germinated spores and small fluorescent puncta in the cytoplasm of appressoria, while mScarlet-*Ch*Snc1 also labelled the appressorium plasma membrane. Fluorescence and DIC channels are presented. Scale bars = 5 μm.

## Tables

**TABLE S1: All proteins detected by mass spectrometry associated with trypsin-treated, density-purified *C. higginsianum* EVs** (a) All detected proteins associated with low density *C. higginsianum* EVs. (b) All detected proteins associated with high density *C. higginsianum* EVs.

(Excel file included separately)

**TABLE S2: *C. higginsianum* EV proteomes (2/4 replicates, q-value ≤ 0.01)** (a) Proteins associated with low density *C. higginsianum* EVs detected in 2/4 replicates with a q-value ≤ 0.01. (b) Proteins associated with high density *C. higginsianum* EVs detected in 2/4 replicates with a q-value ≤ 0.01.

(Excel file included separately)

**TABLE S3:** Secondary metabolism-related proteins associated with *C. higginsianum* EVs

(Excel file included separately)

**TABLE S4: Gene Ontology (GO) Term Enrichment for proteins associated with *C. higginsianum* EVs** (a) GO term enrichment for proteins associated with low density *C. higginsianum* EVs (b) GO term enrichment for proteins associated with high density *C. higginsianum* EVs (c) GO term enrichment for proteins unique to low density *C. higginsianum* EVs (d) GO term enrichment for proteins unique to high density *C. higginsianum* EVs. All proteins analyzed were detected in 2/4 replicates and had a q-value ≤ 0.01.

(Excel file included separately)

**TABLE S5:**
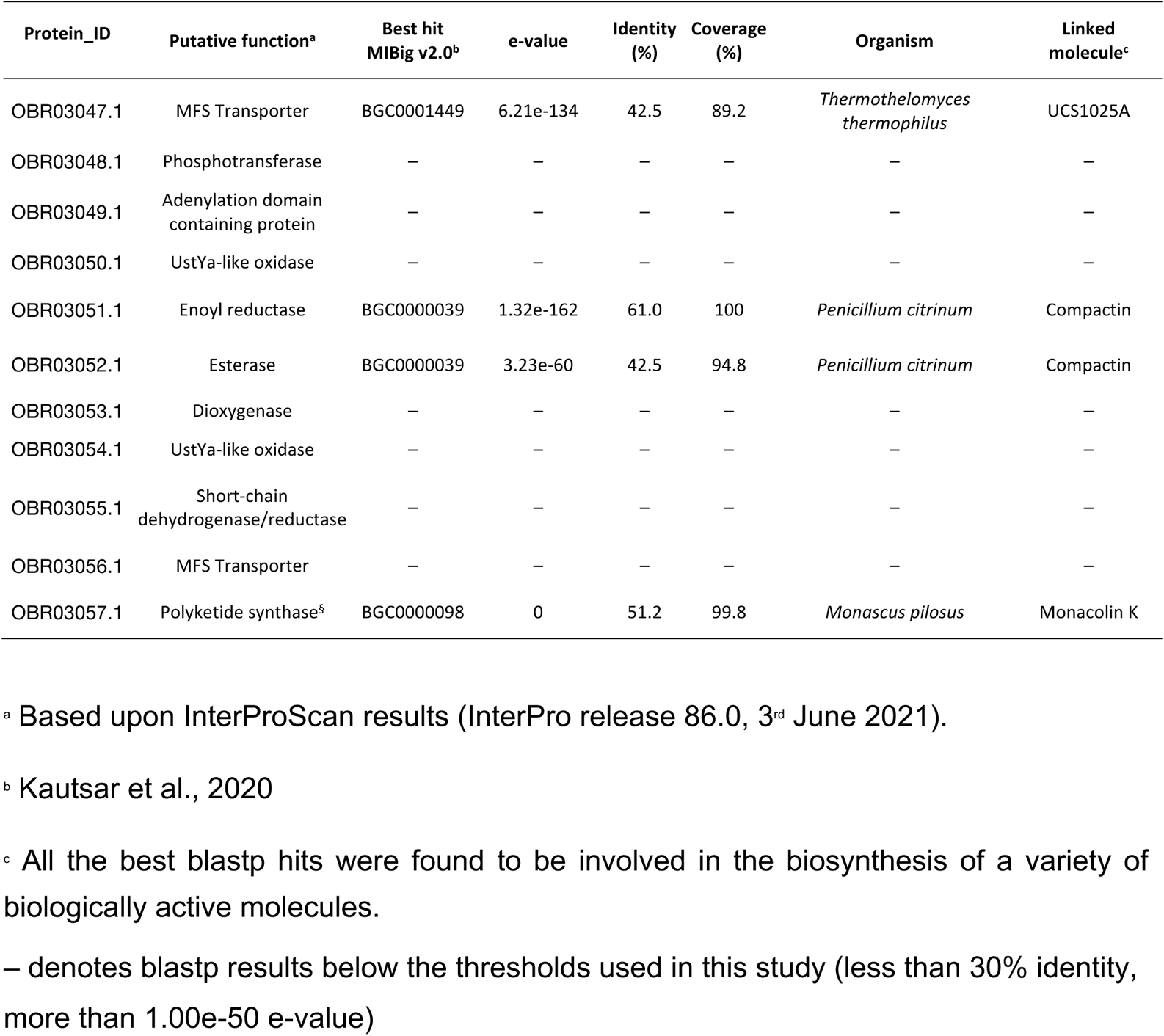
Best blastp hits from MIBiG database of BGCs linked to their biosynthetic product for the BGC21 of Colletotrichum higginsianum.

**TABLE S6:**
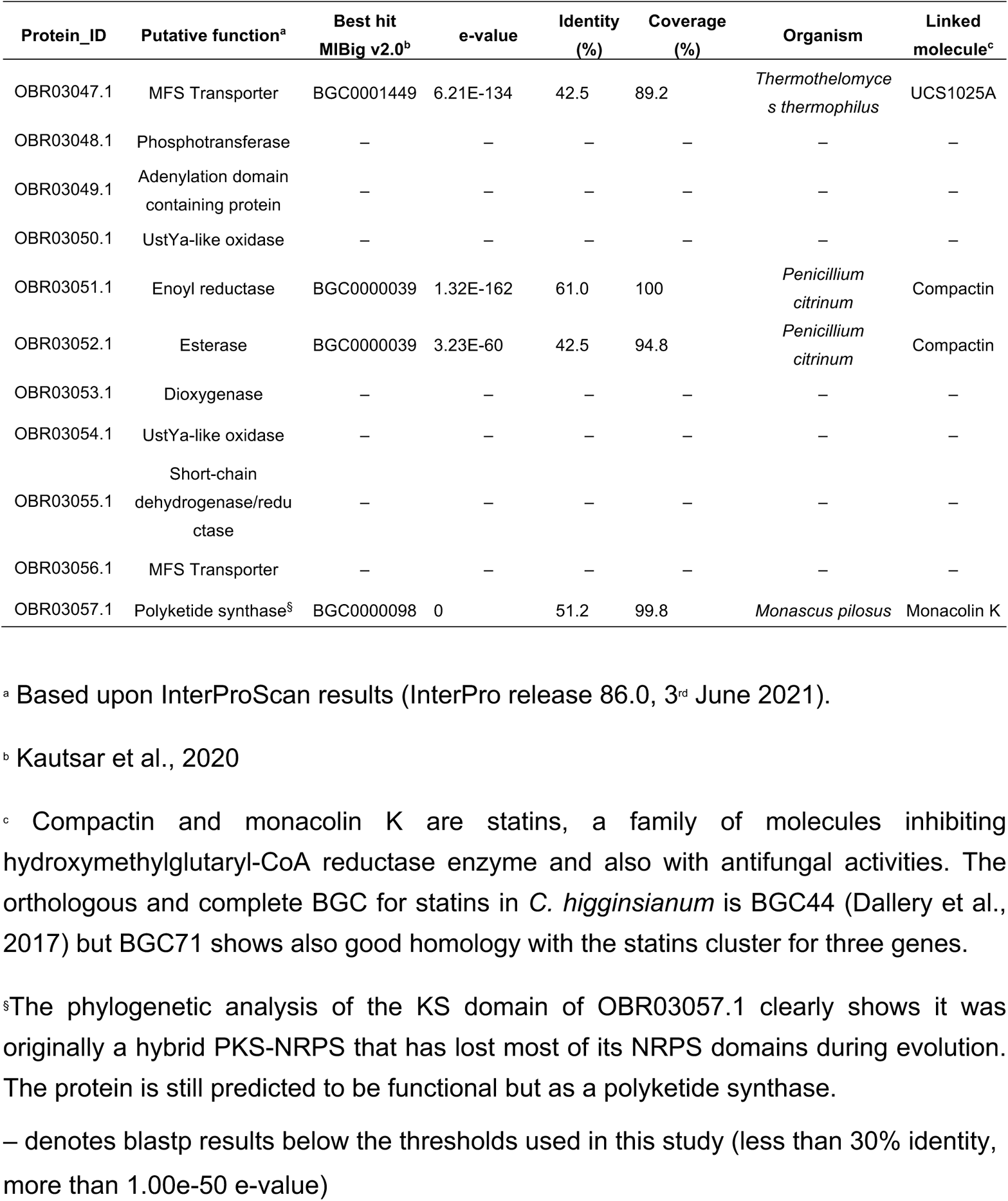
Best blastp hits from MIBiG database of BGCs linked to their biosynthetic product for the BGC71 of Colletotrichum higginsianum.

**TABLE S7:**
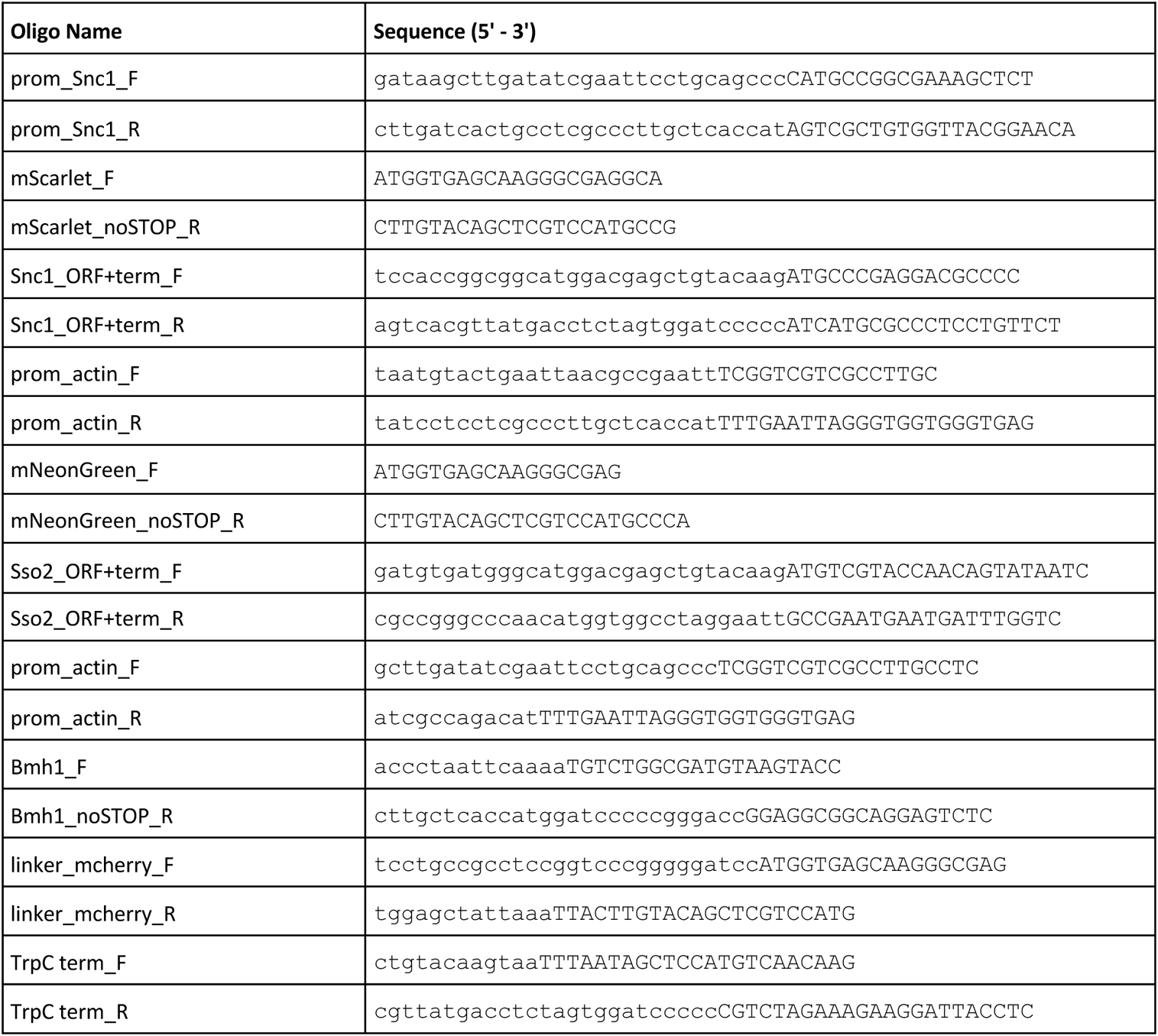
List of primers used for cloning.

